# Combinatorial circuit dynamics orchestrate flexible motor patterns in *Drosophila*

**DOI:** 10.1101/2022.12.14.520499

**Authors:** Hiroshi M. Shiozaki, Kaiyu Wang, Joshua L. Lillvis, Min Xu, Barry J. Dickson, David L. Stern

**Author notes:** Correspondence (H.M.S.), (D.L.S.).

## Abstract

Motor systems flexibly implement diverse motor programs to pattern behavioral sequences, yet their neural underpinnings remain unclear. Here, we investigated the neural circuit mechanisms of flexible courtship behavior in *Drosophila*. Courting males alternately produce two types of courtship song. By recording calcium signals in the ventral nerve cord (VNC) in behaving flies, we found that different songs are produced by activating overlapping neural populations with distinct motor functions in a combinatorial manner. Recordings from the brain suggest that song is driven by two descending pathways – one defines when to sing and the other specifies what song to sing. Connectomic analysis reveals that these “when” and “what” descending pathways provide structured input to VNC neurons with different motor functions. These results suggest that dynamic changes in the activation patterns of descending pathways drive different combinations of motor modules, thereby flexibly switching between different motor actions.

## Introduction

Animals flexibly switch between different actions to adapt to a changing environment. A common mechanism to produce switching actions is to activate separate neural populations, each dedicated to one action. For example, during locomotion, flexor and extensor muscles in the vertebrate limb alternately contract through activation of different neural populations in the spinal cord^1, 2^. This type of motor control often underlies body movements that involve contractions of distinct populations of muscles. Diverse actions can also be produced by differentially contracting overlapping sets of muscles, as seen in the respiratory system^3^, yet little is known about how these actions are controlled by the motor system.

During courtship, *Drosophila melanogaster* males vibrate their wings in specific patterns to produce acoustic communication signals called courtship song. The song is composed of two primary types, pulse and sine^4^ (Figure 1A), which influence females’ receptivity to males in different ways^5, 6^. Males dynamically switch between pulse and sine song depending on sensory feedback from females^7^ as well as based on internal states^8^. The sounds of song are shaped by wing control muscles^9^, which induce rapid changes in wing motion through coordinated firing^10^. Most of the control muscles are active during pulse song, whereas a subset of these active muscles becomes silent during sine song^10, 11^. Thus, the two song patterns are produced by recruiting overlapping, rather than distinct, sets of wing control muscles.

**Figure 1.**
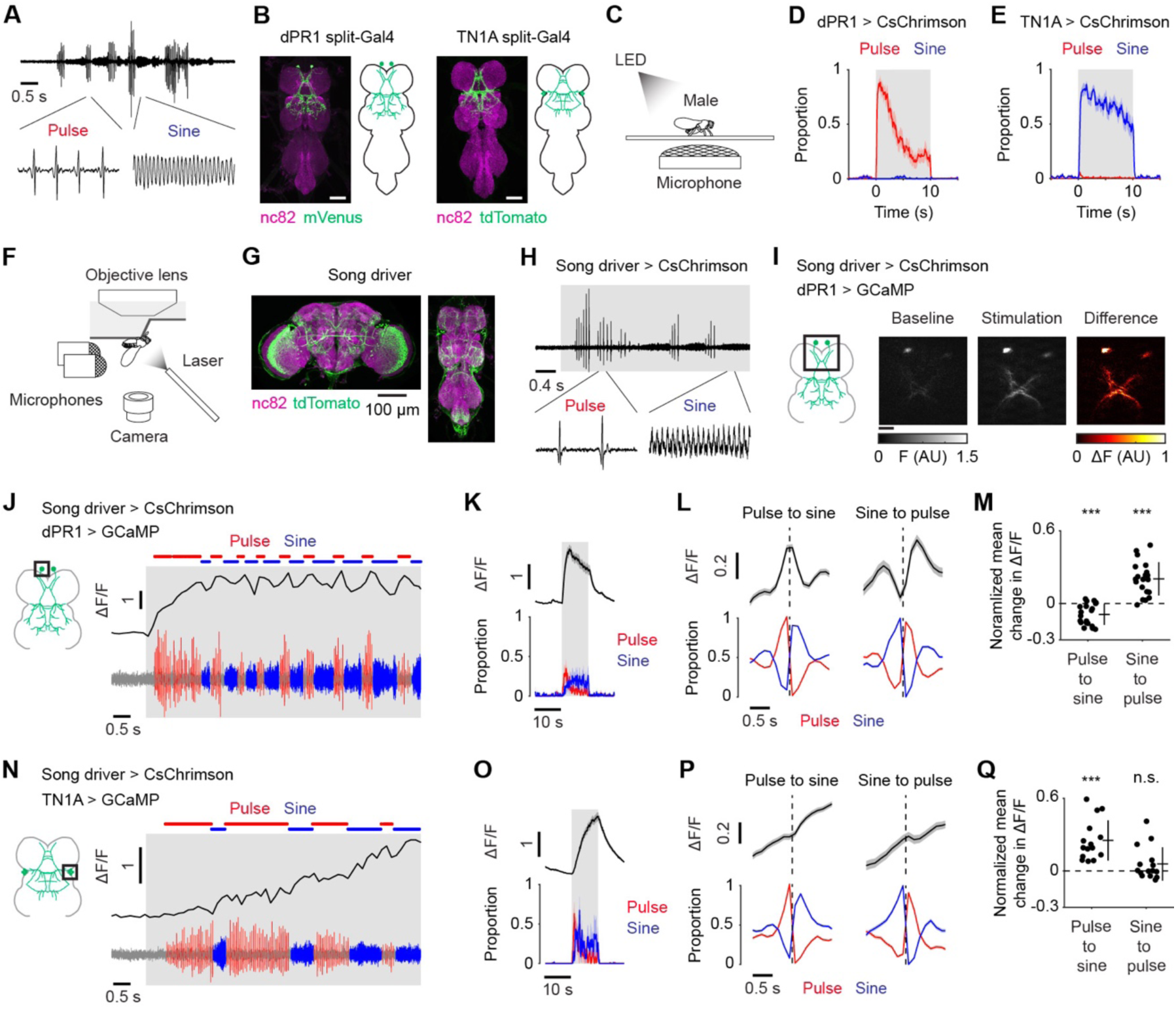
Flexible song production involves combinatorial activation of dPR1 and TN1A neurons. (A) A recording of the sound of natural *Drosophila melanogaster* courtship song. (B) Expression patterns of the dPR1 split-Gal4 (left) and the TN1A split-Gal4 (right) in the VNC. Scale bars, 50 μm. (C) Schematic of the optogenetic activation experiment. (D) Time course of the proportions of pulse (red) and sine (blue) song induced by LED stimulation of dPR1 at an irradiance of 15.9 μW/mm^2^. Data are represented as mean ± SEM across flies (N = 12 flies). (E) Same as (D) but for TN1A (N = 12 flies). (F) Schematic of calcium imaging during fly singing. (G) Expression pattern of the driver line used to induce pulse and sine song with optogenetic activation. (H) Example of the sounds induced by optogenetic activation of the song driver in a fly attached to the recording plate. No thoracic dissection was made prior to this recording. The shaded area represents the period when laser stimulation was applied. (I) Example frames during recording from dPR1. Scale bar, 20 μm. (J) Example ΔF/F trace of a single dPR1 neuron (top) together with the simultaneously recorded sound (bottom). (K) Time course of ΔF/F recorded from dPR1 (top) and the proportions of pulse and sine song (bottom) during laser stimulation at an irradiance of 9.8 μW/mm^2^. Data are represented as mean ± SEM across neurons (top; N = 18 neurons in 9 flies) and flies (bottom; N = 9 files). (L) Time course of dPR1 ΔF/F (top) and song (bottom) around song-type transitions. Dashed vertical lines represent the timing of transitions. Data are from 20 neurons in 10 flies and represented as mean ± SEM across transitions for both ΔF/F and song (N = 2184 and 858 events for pulse-to-sine and sine-to-pulse transitions, respectively). (M) The mean change in ΔF/F after song-type transitions relative to ΔF/F before the transitions (see STAR Methods for detail) for dPR1. Each dot represents a neuron. Lines represent mean ± SD across neurons (N = 20 neurons in 10 flies). ***P < 0.001/2, one-sample t test with Bonferroni correction, the denominator represents the number of comparisons. (N–Q) Same as (J–M) but for TN1A. (O) N = 15 neurons in 4 flies. (P) N = 967 (pulse-to-sine) and 444 (sine-to-pulse) transitions from 15 neurons in 4 flies. (Q) N = 15 neurons in 4 flies. See also Figure S1.

Wing motor neurons extend dendrites into the ventral nerve cord (VNC)^9, 11–13^ where they receive input from thoracic interneurons that, in turn, integrate song promoting descending commands from the brain^4, 8, 14–19^. Activity of VNC interneurons is sufficient to generate song^16, 20^, indicating that the song motor circuit resides in the VNC. Neurons that express the sex determination genes *fruitless* (*fru*) and/or *doublesex* (*dsx*) are involved in song production^19, 21, 22^., and some of these neurons appear to contribute differentially to pulse and sine song. For example, silencing the activity of dPR1 neurons impairs pulse song production^19, 23^, whereas silencing TN1A neurons reduces sine but not pulse song production^21, 23^. However, it remains unclear whether and how the activity of these neurons coordinates flexible song production because no existing preparation allows the recording of neural activity in the VNC during singing.

Here, we developed a two-photon calcium imaging assay for recording signals from neurons in the song motor circuits while flies sang alternate bouts of pulse and sine song. We found that dPR1 neurons are selectively active during pulse, whereas TN1A neurons are active during both pulse and sine song. Recordings from other song-related neurons in the VNC suggest that the combinatorial activity patterns displayed by dPR1 and TN1A neurons is a general feature of the song circuit in the VNC. Connectomic analysis showed that dPR1 and TN1A neurons belong to distinct networks that receive differential input from parallel descending pathways. Imaging the activity of descending neurons that synapse onto dPR1 and TN1A neurons suggests that one descending pathway specifies production of song whereas another pathway specifies the song type (pulse versus sine). Together, our results suggest that descending commands drive combinatorial activation of distinct motor modules to flexibly produce different types of acoustic communication signals in *D. melanogaster*.

## Results

### Flexible song production involves combinatorial activation of dPR1 and TN1A neurons

To study the mechanisms of flexible song production, we built driver lines for dPR1 and TN1A using the split-Gal4 method^24, 25^. Both dPR1 and TN1A neurons express the sex determination gene *dsx*^19, 21^. The dPR1 driver line cleanly labels a pair of VNC neurons (Figures 1B, S1A, and S2B) whose cell body locations and innervation patterns match those of dPR1^19^. As expected, these labeled neurons express *dsx* (Figure S1C). The driver line for TN1A was constructed using *dsx*-DBD as a hemi-driver and thus labels a subset of *dsx*-expressing neurons. This line labels approximately four VNC neurons per hemisphere whose morphology matches that of TN1A^21^ (Figures 1B, S1A, S1D, and S1E).

dPR1 and TN1A were previously implicated in pulse and sine song production, respectively^15, 19, 21^. However, due to limitations in available reagents, it has remained unclear how specifically these neurons contribute to each song type. To address this issue, we expressed the light-gated cation channel CsChrimson^26^ in dPR1 or TN1A neurons and applied activation light in solitary males placed in song recording chambers (Figure 1C). Optogenetic activation of dPR1 acutely elicited pulse but not sine song over a range of stimulation intensities, whereas activation of TN1A induced sine but not pulse song^23^ (Figures 1D, 1E, and S1F–J). Thus, dPR1 and TN1A are capable of inducing pulse and sine song, respectively.

We next examined the activity of dPR1 and TN1A neurons during pulse and sine song. There has been no preparation that allows recording the activity of VNC neurons in singing flies. This partly stems from two technical challenges. First, wing vibration is dependent on muscle-induced oscillatory movements of the thorax^27^ and thus attaching the thorax, which houses the VNC, to the recording apparatus often disturbs these oscillations. Second, wing muscles are adjacent to the VNC and are easily damaged during dissection required to image the VNC. To overcome these challenges, we glued parts of the legs, instead of the thorax, to the recording plate and dissected the ventral side of the thorax where wing muscles are absent (Figure 1F). This novel assay provides optical access to VNC neurons while allowing approximately normal wing vibrations for song production.

Song was induced by optogenetic activation of neurons labeled by a LexA line (R22D03-LexA) that we term the “song driver” (Figure 1G). This driver line labels multiple neurons including the male specific brain neurons P1, which integrate multisensory cues and promote the initiation and persistence of courtship behavior^14, 17, 19, 28–32^ (Figure S1K). Optogenetic activation of the neurons labeled by the song driver in tethered males induced alternating bouts of pulse and sine song as in natural singing, even though the activation light was maintained at constant power (Figures 1H, S1L, and S1M). The bout durations of the induced pulse and sine song were similar to those produced during normal courtship in freely moving flies (Figure S1N). Removal of the head eliminated the production of song during optogenetic stimulation (Figures S1M and S1O). This effect was not caused by decapitation-induced physical damage because optogenetic activation of pIP10, the song promoting descending neuron^4, 8, 14–19^, in the same preparation elicited song in decapitated flies (Figures S1P and S1Q). These results indicate that neurons labeled by this driver line in the brain (which includes P1 neurons), but not those labeled in the VNC, are required for inducing song by optogenetic stimulation.

To characterize the activity of dPR1 and TN1A during pulse and sine song, we performed two-photon imaging of the genetically encoded calcium indicator jGCaMP7f^33^ expressed in dPR1 or TN1A neurons with the split-Gal4 lines while inducing song by optogenetic stimulation of the song driver. We extracted the calcium signals of individual neurons by imaging the cell bodies. Optogenetic stimulation acutely increased calcium signals of dPR1 neurons as well as the amount of song, consistent with a role of dPR1 in song production (Figures 1I–K, and S1R–U; Video S1). Notably, over the course of constant optogenetic stimulation, the dPR1 calcium signals repeatedly increased and decreased as the fly alternated between song types (Figure 1J). The calcium signal increased during transitions from sine to pulse and decreased during transitions from pulse to sine (Figures 1J, 1L, and 1M). The preference for pulse is consistent with the observation that optogenetic activation of dPR1 promotes pulse song (Figure 1D).

Calcium dynamics measured at the soma are known to be slower than those at the neuropil^34^. Indeed, dPR1 signals measured at the neuropil showed more rapid modulation than somatic signals (Figures S1V–X). The decay rate of the neuropil signal during sine song (Figure S1V) was comparable to the decay kinetics of jGCaMP7f in flies^33^. These results suggest that dPR1 neurons are active during pulse song but not during sine song.

Similar to the observations we made of dPR1, calcium signals of individual TN1A neurons increased in response to optogenetic stimulation of the song driver (Figures 1N, 1O, and S1Y–BB; Video S2). Consistent with the observation that optogenetic activation of TN1A promotes sine (Figure 1E), TN1A calcium signals increased when the song type switched from pulse to sine (Figures 1N, 1P, and 1Q). However, TN1A calcium signals did not decrease when the song type switched from sine to pulse, but instead remained unchanged (Figures 1N, 1P, and 1Q). This pattern contrasts with the dPR1 calcium signals, which rapidly decreased after the transition to the non-preferred song type (Figure 1L). The different calcium signal patterns between dPR1 and TN1A did not result from slower kinetics of calcium at the cell body than in the neuropil, because TN1A calcium signals measured at the neuropil decreased only slightly when the song type switched from sine to pulse (Figures S1CC–EE). These results suggest that TN1A neurons are active during both pulse and sine song with a preference for sine.

The tethered flies occasionally produced pulse song outside of the optogenetic stimulation periods, as observed during intermittent stimulation of P1 neurons in freely moving isolated male flies^14, 17^, which allowed us to determine whether the activity patterns we had observed for dPR1 and TN1A during pulse song were dependent on activation of the song driver. We found that both dPR1 and TN1A showed increased calcium signals when flies sang pulse song outside of the optogenetic stimulation periods (Figures S1FF and S1GG). This result further supports the idea that pulse song involves activation of both dPR1 and TN1A.

As an independent test of pulse-related activity, we recorded calcium signals from dPR1 and TN1A while pulse song was induced by activating pIP10, a pair of song promoting descending neurons^4, 8, 14–19^. We expressed CsChrimson in pIP10 using a new split-LexA line that cleanly labels pIP10 (Figures 2A and S2A) while driving the expression of jGCaMP7s in dPR1 or TN1A. Optogenetic activation with the split-LexA line induced predominantly pulse song in freely behaving flies (Figures 2B, S2B, and S2C) as well as during calcium imaging (Figures S2E and S2G). Both dPR1 and TN1A showed increased calcium signals when flies sang pulse song (Figures 2C–H and S2D–G. Taken together, our data suggest that flexible song production involves combinatorial activation of dPR1 and TN1A—both cell types are active during pulse song while only TN1A neurons are active during sine song.

**Figure 2.**
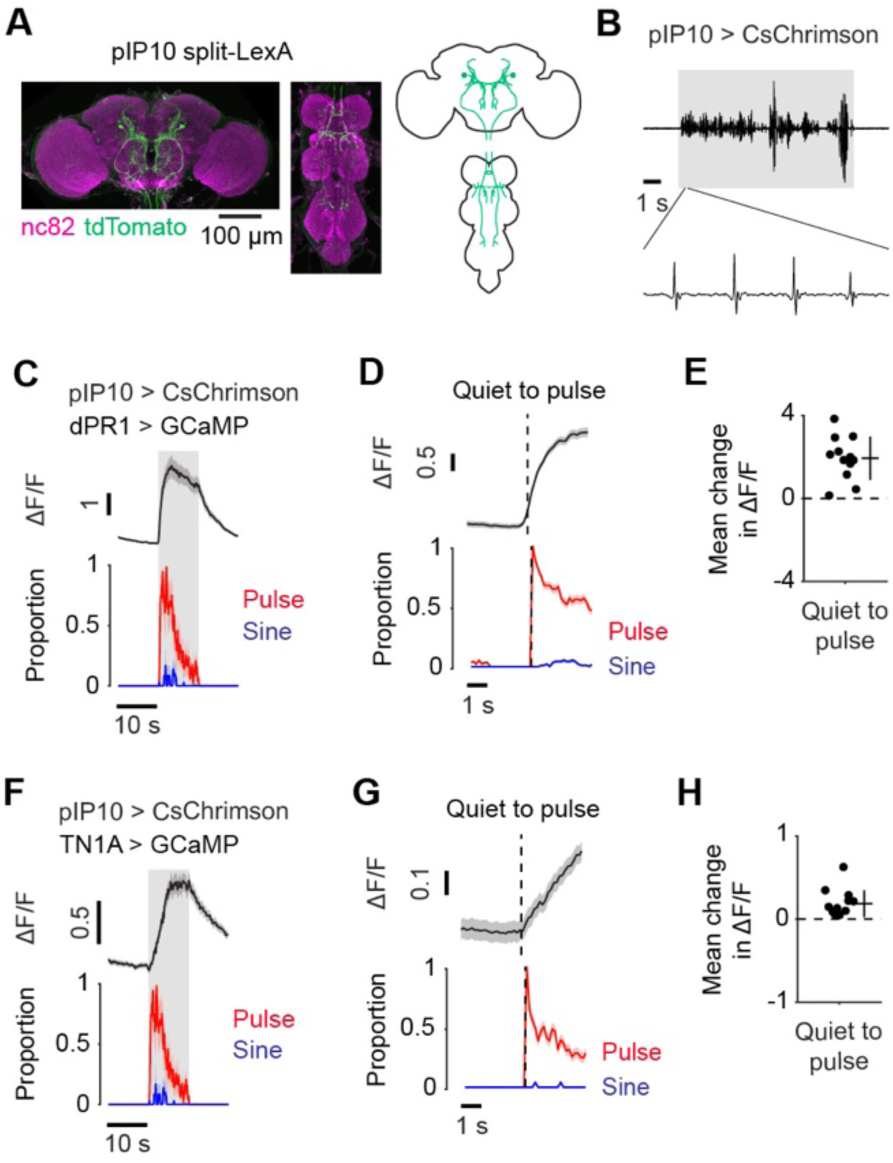
dPR1 and TN1A are activated during pulse song induced by stimulation of the descending neurons pIP10. (A) Expression pattern of the split-LexA driver for pIP10. (B) Example of the sound induced by optogenetic activation of pIP10 in the single fly optogenetic activation experiment. The shaded area represents the period of LED stimulation. (C) Time course of ΔF/F recorded from dPR1 and the proportions of song for the laser stimulation at the irradiance of 156.2 μW/mm^2^. jGCaMP7s was expressed with *dsx*-Gal4 and calcium signals were recorded from dPR1 cell bodies. Data are represented as mean ± SEM across neurons (top; N = 12 neurons in 6 flies) and flies (bottom; N = 6 files). (D) Time course of dPR1 ΔF/F and song around transitions from quiet to pulse song production. Dashed vertical lines represent the transition time. Data are from 12 neurons in 6 flies and represented as mean ± SEM across transitions for both ΔF/F and song (N = 190 transitions). (E) The mean change in ΔF/F after quiet-to-pulse transitions (see STAR Methods for detail) for dPR1. Each dot represents a neuron. Lines represent mean ± SD across neurons (N = 12 neurons in 6 flies). (F–H) Same as (C–E) but for TN1A. GCaMP was expressed with the TN1A split-Gal4 line. (F) N = 19 neurons in 6 flies. (G) N = 99 transitions from 13 neurons in 4 flies. (H) N = 13 neurons in 4 flies. See also Figure S2.

### TN1 neurons exhibit combinatorial activation during flexible song production

To examine whether the combinatorial activation pattern observed in dPR1 and TN1A is a general property of pulse- and sine-preferring neurons in the VNC, we focused on TN1 neurons. These neurons are male-specific, *dsx*-expressing VNC neurons involved in song production^18, 21, 35^. There are approximately twenty-two TN1 neurons in each hemisphere^35^, which are composed of at least five anatomically defined cell types including TN1A^21^ (Figure 3A). While manipulation of the activity of TN1 neurons primarily influences sine song, a subset of TN1 neurons influences pulse song^18, 21^.

**Figure 3.**
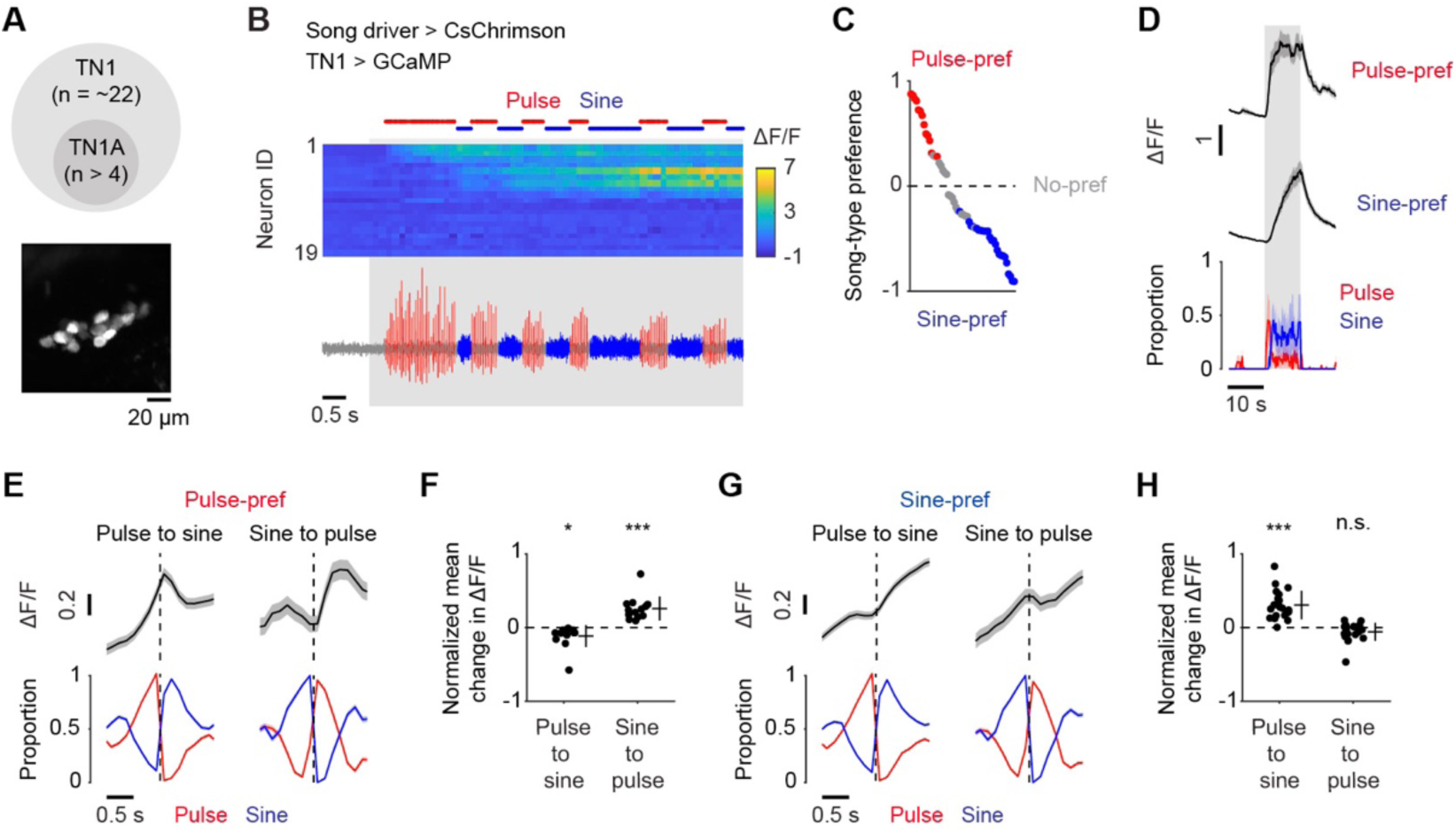
TN1 neurons exhibit combinatorial activation during flexible song production. (A) Top, there are approximately twenty two TN1 neurons in each hemisphere of the VNC^35^. TN1A neurons are a subset of all TN1 neurons. Bottom, example calcium signals from TN1 neurons in one hemisphere *in vivo*. GCaMP was driven by *dsx*-Gal4. The image is a maximum projection of a stack of 10 z-planes. (B) Example ΔF/F of a population of TN1 cell bodies recorded simultaneously in a single individual (top) together with the sound recording (bottom). The song driver was used to express CsChrimson. The shaded area represents the period where laser stimulation was applied. (C) The distribution of the song-type preference (N = 56 neurons in 4 flies), which quantifies how much calcium signals differed between pulse and sine song around song-type transitions (see STAR Methods for detail). Red and blue dots indicate neurons whose calcium signals were significantly higher for pulse and sine, respectively (p < 0.05, permutation test). (D) Time course of ΔF/F recorded from pulse-preferring and sine-preferring TN1 neurons (top) and the proportions of pulse and sine song (bottom) for the laser stimulation at an irradiance of 9.8 μW/mm^2^. Data are represented as mean ± SEM across neurons (top; N = 13 pulse-preferring in 3 flies and 25 sine-preferring neurons in 4 flies) and flies (bottom; N = 4 files). (E) Time course of ΔF/F of pulse-preferring TN1 neurons (top) and song (bottom) around song-type transitions. Dashed vertical lines represent the transition time. Data are from 13 neurons in 3 flies and represented as mean ± SEM across transitions for both ΔF/F and song (N = 825 and 315 events for pulse-to-sine and sine-to-pulse transitions, respectively). (F) The mean change in ΔF/F after song-type transitions relative to ΔF/F before the transitions (see STAR Methods for detail) for pulse-preferring TN1 neurons. Each dot represents a neuron. Lines represent mean ± SD across neurons (N = 13 neurons in 3 flies). (G and H) Same as (E) and (F) but for sine-preferring TN1 neurons. (G) N = 1370 (pulse-to-sine) and 511 (sine-to-pulse) transitions from 25 neurons in 4 flies. (H) N = 21 neurons in 4 flies. Four neurons were excluded from this analysis because the mean ΔF/F before song type transitions was too low (<0.1) to reliably estimate the normalized change in ΔF/F. See also Figure S3.

To characterize the activity of TN1 neurons during song, we recorded calcium signals from a large fraction of individual TN1 neurons in each fly (Figure 3A and 3B) while inducing song by optogenetic stimulation of the song driver. The optogenetic stimulation elicited significant changes in calcium signals (p < 0.05, t-test, see STAR Methods for detail) in 69.1% of the recorded TN1 neurons (56/89 neurons in 4 flies). For each neuron with significant responses, we quantified the selectivity for song type with an index we termed the song-type preference (Figure 3C, S3A, and S3B). Most TN1 neurons showed significant preference for pulse or sine song (38/56 neurons in 4 flies; p < 0.05, permutation test), with most preferring sine song (N = 25/38 neurons in 4 flies) (Figure 3C). The average number of sine-preferring neurons in each fly (N = 6.3 neurons per hemisphere) was larger than the number of neurons labeled by the TN1A split-Gal4 driver (N = 4 neurons per hemisphere), suggesting that the sine-preferring population includes neurons that were not labeled with the TN1A split-Gal4 line discussed earlier.

As observed for dPR1 and TN1A, pulse- and sine-preferring TN1 neurons displayed increased calcium signals during optogenetic stimulation of the song driver (Figure 3D, S3C, and S3C). To examine the pulse- and sine-related activity, we calculated the average calcium signals during song-type transitions for each population. We found that both populations showed increased calcium signals when the song switched to the preferred type (Figure 3E–H). However, pulse-but not sine-preferring neurons showed decreased calcium signals when the song switched to the non-preferred type (Figures 3E–H). This result suggests that—as we observed for dPR1 and TN1A neurons—both pulse- and sine-preferring TN1 neurons are active during pulse song, while only the sine-preferring neurons show high activity levels during sine song.

To further characterize pulse-related activity in TN1 neurons, we drove pulse song by optogenetically activating pIP10, which primarily drives pulse song, while recording calcium signals from TN1 neurons. Optogenetic stimulation of pIP10 induced changes in calcium signals in a large fraction of TN1 neurons (77/83 neurons in 4 flies; p < 0.05, t-test, see STAR Methods for detail), of which virtually all neurons displayed increased calcium signals when the flies produced pulse song (76/77 neurons; Figures S3E–J). This result further supports the idea that pulse song involves the coactivation of pulse-preferring and sine-preferring TN1 neurons during pulse song.

### Combinatorial activation is a general feature in the song motor circuit

To further examine the generality of combinatorial activation during flexible song production, we expressed jGCaMP7f with a pan-neuronal driver and performed volumetric calcium imaging in the wing-related neuropils (Figure 4A)^36^, which are innervated by at least tens of cell types^11, 13, 19, 21, 22, 30, 37^, while inducing song with optogenetic stimulation of the song driver (Figure 4B). Since two-photon microscopy cannot resolve individual neuronal processes within dense projections, the measured calcium signal in each voxel likely reflected the combined activity of multiple neurons. Optogenetic stimulation elicited an increase in calcium signals in a large proportion of the recorded volume (Figure 4C; Video S3). A large fraction of the activated voxels showed comparable calcium signals during pulse and sine song (Figure 4C–E and 4G), suggesting that most of the song-related neurons are similarly active irrespective of song type. We also found that a subset of the activated voxels modulated calcium signals when the fly switched between pulse and sine song (Figure 4C, 4D, 4E, 4G, S4A–C; Videos S3 and S4). Notably, most of these song-type selective voxels preferred pulse song (Figures 4D–H and S4A–C). These observations suggest that the combinatorial activity pattern observed in cell-type specific recording of dPR1 and TN1A neurons is a general property of the song circuit during flexible song production—a population of neurons is active both during pulse and sine song while another population is selectively active during pulse song.

**Figure 4.**
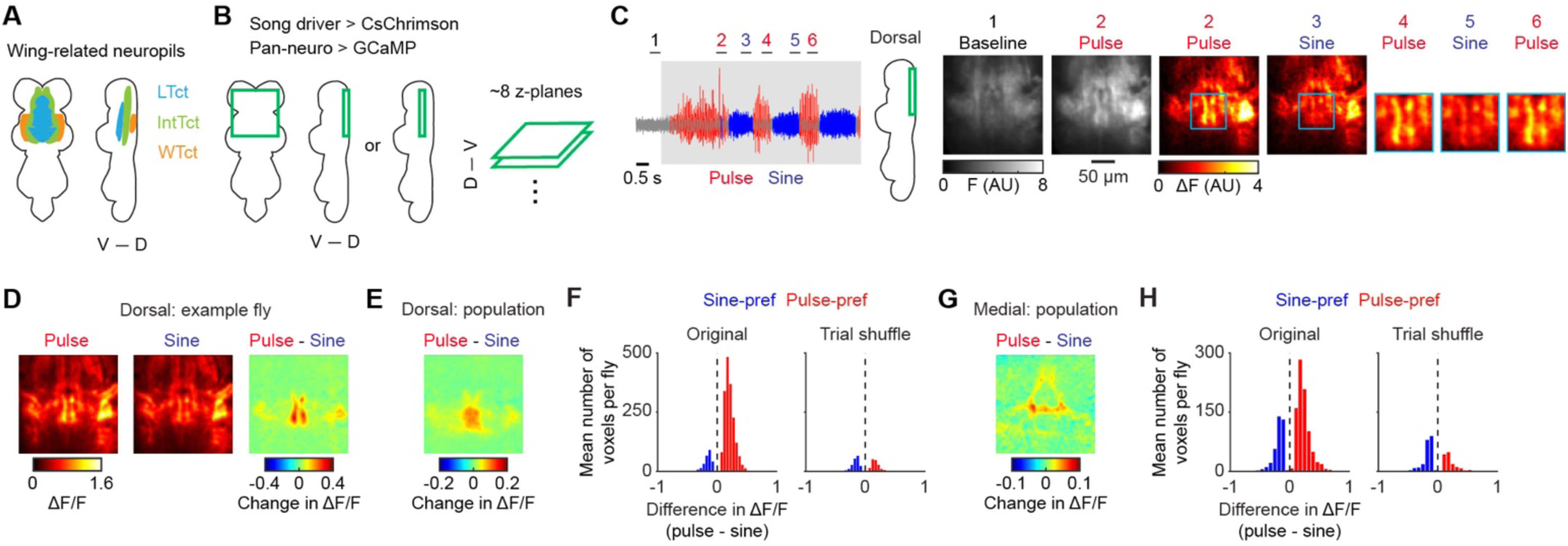
Combinatorial activation is a general feature in the song motor circuit. (A) Schematic of the wing-related neuropils. LTct, lower tectulum. IntTct, intermediate tectulum. WTct, wing tectulum. D, dorsal; V, ventral. (B) Schematic of the region imaged. For each fly, images were obtained from either a dorsal or medial part of the VNC. CsChrimson and GCaMP were expressed using the song and pan-neuronal drivers, respectively. (C) Left, The sound data from an example recording. Bars at the top indicate periods during which example calcium imaging frames, shown on the right, are presented. The shaded area represents the period during which laser stimulation was applied. Right, Example frames. For each period, images were averaged across z-planes and time. For periods 4–6, only part of the images (blue boxes in periods 2 and 3) is shown. (D) The mean ΔF/F during pulse (left), sine (middle), and their difference (right) around song-type transitions for the dorsal part of the VNC in the example fly (N = 100 transitions). Images were averaged across z-planes and time. (E) Population averaged ΔF/F for the difference between pulse and sine song around song-type transitions for the recordings from the dorsal part of the VNC (N = 9 flies). (F) Histogram of the mean difference in ΔF/F between pulse and sine song during song-type transitions for the voxels which changed ΔF/F depending on song type (N = 9 flies; p < 0.001, permutation test). To obtain a histogram expected by chance, the same analysis was conducted after randomizing the relationship between song and calcium signals (“Trial shuffle”, see STAR Methods). (G and H) Same as (E) and (F) but for the recordings from the medial part (N = 3 flies). See also Figure S4.

The song driver might label neurons that are not related to song production, raising the possibility that optogenetic stimulation of these neurons contributed to voxel activation during both pulse and sine song. To test this possibility, we examined the patterns of pan-neuronal calcium signals induced by optogenetic stimulation of the dPR1 or TN1A split-Gal4 drivers (Figure S4D), which cleanly labels these cell types and induces pulse and sine song, respectively (Figure 1B). Stimulation of either cell type led to widespread responses in largely overlapping volumes (Figures S4E and S4F), as expected from the neurons active during both pulse and sine song. Yet, the activation patterns induced by the stimulation of these cell types were not identical. For example, dPR1 but not TN1A activation induced strong responses along the anterior-posterior axis around the midline in the wing tectulum (Figures S4E and S4F), in the same region where pulse-preferring voxels were observed during stimulation of the song driver (Figure 4E and S4B). These differential response patterns were not explained by calcium signals of the neurons experiencing optogenetic stimulation because dPR1 and TN1A neurons innervate highly overlapping regions (Figures S4G and S4H). These results support the idea that flexible song production involves two neural populations, one active during both pulse and sine song and the other active selectively during pulse song.

### Song type selective signals in the motor circuit occur in a genetically defined population of neurons

During pan-neuronal imaging, the pulse-preferring were localized to a region called the mesothoracic triangle in the medial VNC^22^ and the medial and lateral parts of the dorsal VNC (Figures 4E and 4G). We noticed that this distribution is similar to the innervation pattern of neurons expressing *fruitless* (*fru*)^19, 22^ (Figure 5A), a sex determination gene that regulates sexually dimorphic behaviors including courtship^38^. There are approximately 500 *fru*-expressing neurons in the male VNC, comprising at least 35 morphologically distinct cell types^19, 22^. A subset of these cell types, including dPR1, are known to innervate a VNC region overlapping the pulse-preferring voxels^19, 22^. Thus, these neurons might be the source of the pulse-selective signals observed in the pan-neuronal recording.

**Figure 5.**
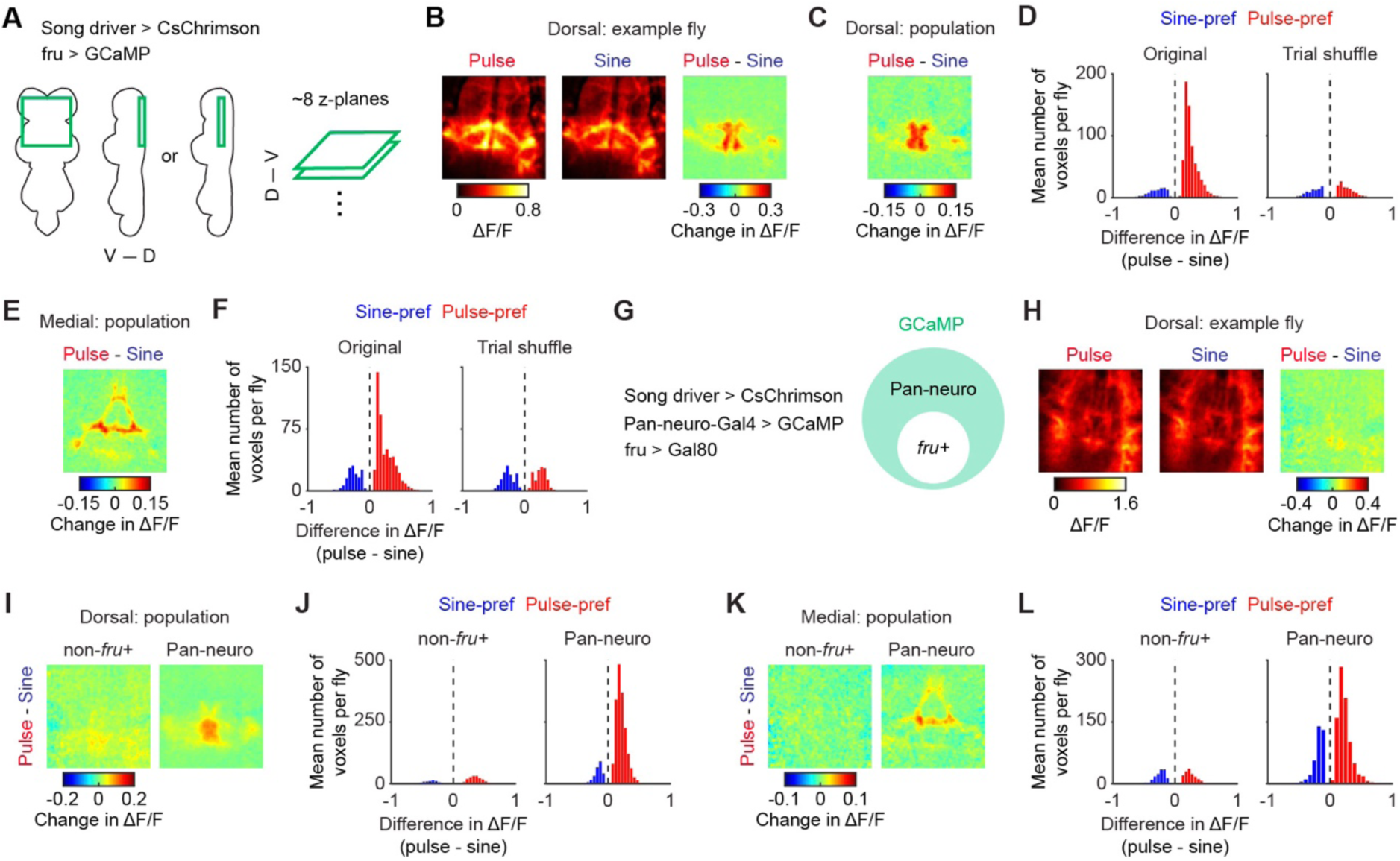
Song type selective signals in the motor circuit are carried by a genetically defined population of neurons. (A) Schematic of the imaged region. For each fly, images were obtained from either dorsal or medial regions of the VNC. CsChrimson and GCaMP were expressed using the song driver and *fru*-Gal4, respectively. D, dorsal; V, ventral. (B) The mean ΔF/F during pulse (left), sine (middle), and their difference (right) around song-type transitions for the dorsal part of the VNC in an example fly (N = 202 transitions). Images were averaged across z-planes and time. (C) Population averaged ΔF/F for the difference between pulse and sine song around song-type transitions for the recordings from the dorsal part of the VNC (N = 6 flies). (D) Histogram of the mean difference in ΔF/F between pulse and sine song during song-type transitions for the voxels which changed ΔF/F depending on song type (p < 0.001, permutation test). Data of the recordings from the dorsal part of the VNC were combined (N = 6 flies). A histogram expected by chance was obtained using a randomization method (“Trial shuffle”, see STAR Methods). (E and F) Same as (C) and (D) but for the recordings from the medial part (N = 5 flies). (G–L) Calcium imaging of non-*fru*-expressing neurons. (G) GCaMP was expressed using the pan-neuronal driver while its expression was suppressed in *fru*-expressing neurons. (H) Same as (B) but for non-*fru*-expressing neurons (N = 47 transitions). (I) Population averaged ΔF/F for the difference between pulse and sine song around song-type transitions for the recordings from the dorsal part of the VNC for non-*fru*-expressing neurons (Left, N = 5 flies). The pan-neuronal imaging result described in Figure 4E is also shown for comparison (Right). (J) Histograms of the mean difference in ΔF/F between pulse and sine song during song-type transitions for voxels that changed ΔF/F depending on song-type (p < 0.001, permutation test). Left, non-*fru*-expressing neurons (N = in 5 flies). Right, the pan-neuronal imaging result shown in Figure 4F. (K and L) Same as (I) and (J) but for the recordings from the medial part (N = 3 flies). See also Figure S5.

To test this possibility, we expressed jGCaMP7f using *fru*-Gal4^39^ and recorded calcium signals in the wing-related neuropils while inducing song by optogenetic activation of the song driver (Figure 5A). We found that some voxels showed similar patterns activation during pulse and sine song (Figure 5B, 5C, 5E, and S5B–D), suggesting that a population of *fru* neurons are similarly active irrespective of song type. We also found that some voxels modulated calcium signals depending on song type. Most of these voxels showed stronger signals during pulse than sine song (Figures 5B–F and S5B–D), indicating that some *fru* neurons are more active during pulse than sine song. The locations of these pulse-preferring voxels (Figures 5C and 5E) matched those detected with pan-neuronal imaging (Figures 4E and 4G). Thus, the activity of a population of *fru*-expressing neurons contributes to the pulse-selective signals observed during pan-neuronal imaging.

To examine the extent to which *fru*-expressing neurons account for the pulse-selective signals observed in pan-neuronal imaging, we took an intersectional genetic approach to express GCaMP in all non-*fru* neurons. We used a pan-neuronal driver to express Gal4, which drove the expression of jGCaMP7f, while suppressing jGCaMP7f expression in *fru* neurons with the Gal4 inhibitor, Gal80^22^ (Figure 5G). A large proportion of the imaged volume was similarly activated during pulse and sine song (Figure 5H, 5I, 5K, and S5E–G), as in the pan-neuronal imaging (Figure 5B, 5C, 5E, and S5B–D). However, we observed a profound reduction in the voxels selective for song type when jGCaMP7f expression was suppressed in *fru* neurons (Figures 5H–L and S5E–G). Notably, *fru* neurons comprise only a few percent of the neurons in the VNC^22, 40^, indicating that these neurons have a disproportional impact on the pulse-selective signals. These results suggest that *fru*-expressing neurons play a key role in flexible song production.

### Flexible song production involves combinatorial activation of parallel descending pathways

We next characterized descending signals provided from the brain to the VNC during song. Two types of descending neurons, pIP10 and pMP2, extensively innervate the wing-related neuropils^16, 19, 22^. Optogenetic activation of pIP10 induces both pulse and sine song^16, 23^, whereas pMP2 activation leads to pulse but not sine song^23^. These findings suggest that pIP10 and pMP2 play distinct roles in song production. However, it remains unclear whether and how these neurons are activated to pattern song sequences.

To examine activity patterns of pIP10 and pMP2 during song, we expressed jGCaMP7f in one cell type at a time with split-Gal4 drivers^16, 23^ and recorded calcium signals in the brain of the fly standing on a ball (Figures 6A and 6B). Song was induced by optogenetic activation of the song driver. We found that pIP10 showed increased calcium signals during optogenetic activation as expected for a role in song production (Figures 6C, 6D, S6A, and S6B). These neurons displayed similar levels of calcium signals during pulse and sine song, with a weak preference for pulse (Figures 6C, 6E, and 6F). These results suggest that pIP10 is active during both pulse and sine song.

**Figure 6.**
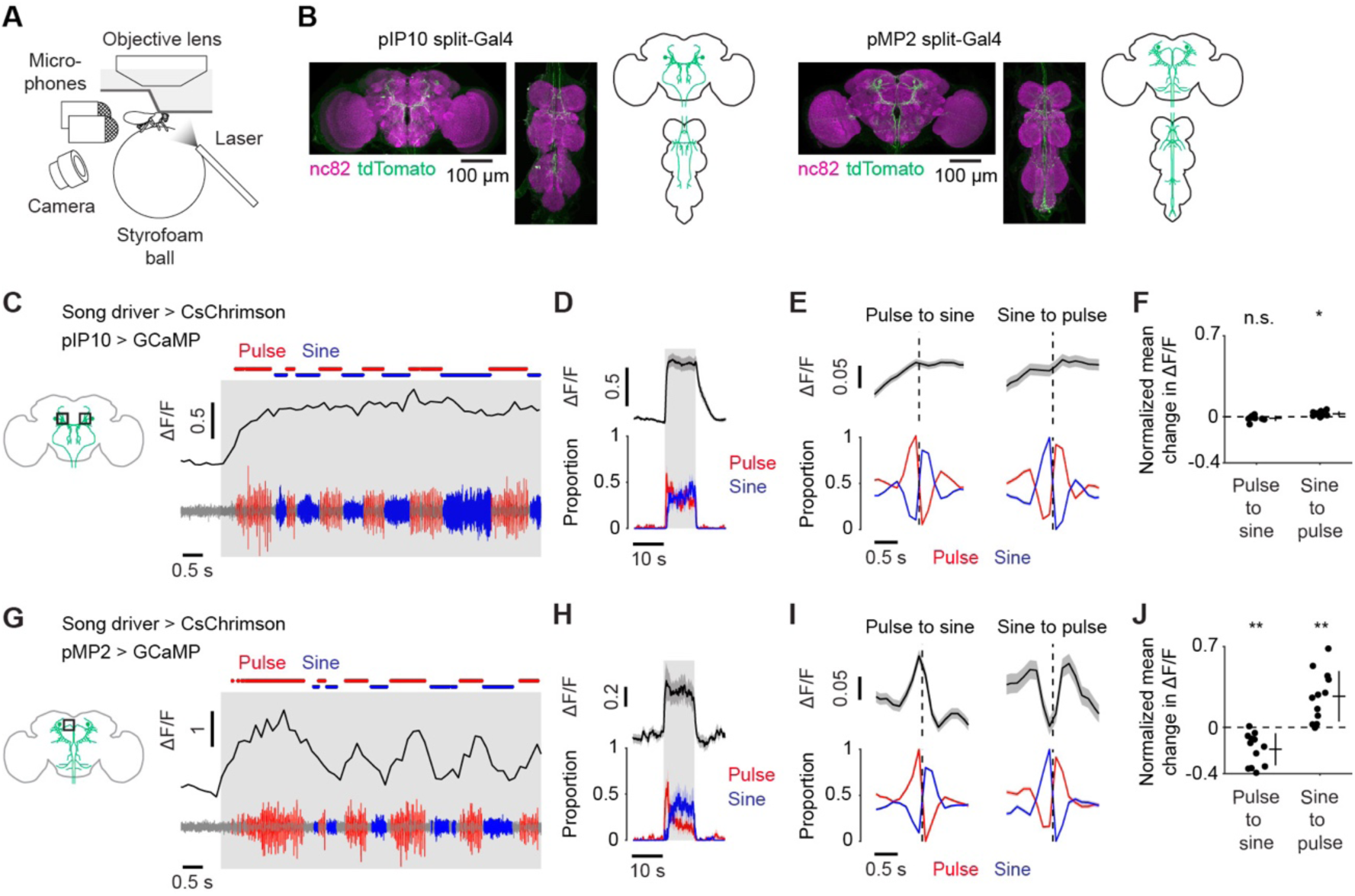
Flexible song production involves combinatorial activation of parallel descending pathways. (A) Schematic of calcium imaging in the brain during fly singing. (B) Expression patterns of the pIP10 split-Gal4 (left) and the pMP2 split-Gal4 (right). The confocal stacks for the pIP10 split-Gal4 were from Ding, Lillvis et al. (2019)^16^. (C) Example ΔF/F trace of pIP10 neurons (top) together with the simultaneously recorded sound (bottom). Calcium signals reflect combined activity of the left and right pIP10 neurons. (D) Time course of ΔF/F recorded from pIP10 (top) and the proportions of pulse and sine song (bottom) during laser stimulation at an irradiance of 575.6 μW/mm^2^. Data are represented as mean ± SEM across neurons (N = 9 flies). (E) Time course of pIP10 ΔF/F (top) and song (bottom) around song-type transitions. Dashed vertical lines represent the timing of transitions. Data are from 9 flies and represented as mean ± SEM across transitions for both ΔF/F and song (N = 2,827 and 1,145 events for pulse-to-sine and sine-to-pulse transitions, respectively). (F) The mean change in ΔF/F after song-type transitions relative to ΔF/F before the transitions (see STAR Methods for detail) for pIP10. Each dot represents a neuron. Lines represent mean ± SD across neurons (N = 9 flies). *P < 0.05/2, one-sample t test with Bonferroni correction, the denominator represents the number of comparisons. (G–J) Same as (C–F) but for pMP2. Calcium signals were recorded from neurites of individual pMP2 neurons. (H) N = 12 neurons in 7 flies. (I) N = 1,487 (pulse-to-sine) and 524 (sine-to-pulse) transitions from 12 neurons in 7 flies. (J) N = 12 neurons in 7 flies. See also Figure S6.

Similar to pIP10, pMP2 showed increased calcium signals during optogenetic activation (Figures 6G, 6H, S6C, and S6D). However, in contrast to pIP10, pMP2 signals were strongly selective for song type—calcium signals increased during pulse song and decreased during sine song (Figures 6G, 6I, and 6J). The pattern of calcium signal modulation was similar to the pattern observed for dPR1 (Figure 1). These results suggest that pMP2 is active during pulse but not sine song. Taken together, parallel descending pathways exhibit combinatorial activation patterns similar to those we observed for the VNC. One pathway, pIP10, is active irrespective of the song type the fly sings, whereas the other pathway, which includes pMP2, is active only during pulse song. These observations suggest that parallel descending pathways independently control the timing and content of the song through the motor circuits in the VNC, which we further characterized by examining synaptic connectivity amongst these neurons.

### Connectome analysis reveals the neural circuit architecture underlying flexible song production

To examine how combinatorial activity patterns in the VNC are driven by descending input, we analyzed the synaptic connectivity among pIP10, pMP2, dPR1, and TN1A neurons using the connectome of the male VNC^41^. All of these neurons are cholinergic^23^ and are likely to be excitatory. We identified these neurons in the connectome dataset based on morphological characteristics. We found only two neurons, one per hemisphere, for pIP10, pMP2, and dPR1 as reported previously^16, 19, 22, 23^ (Figure 7A). There were 24 neurons whose morphology matched TN1A^21^ (Figure 7A).

**Figure 7.**
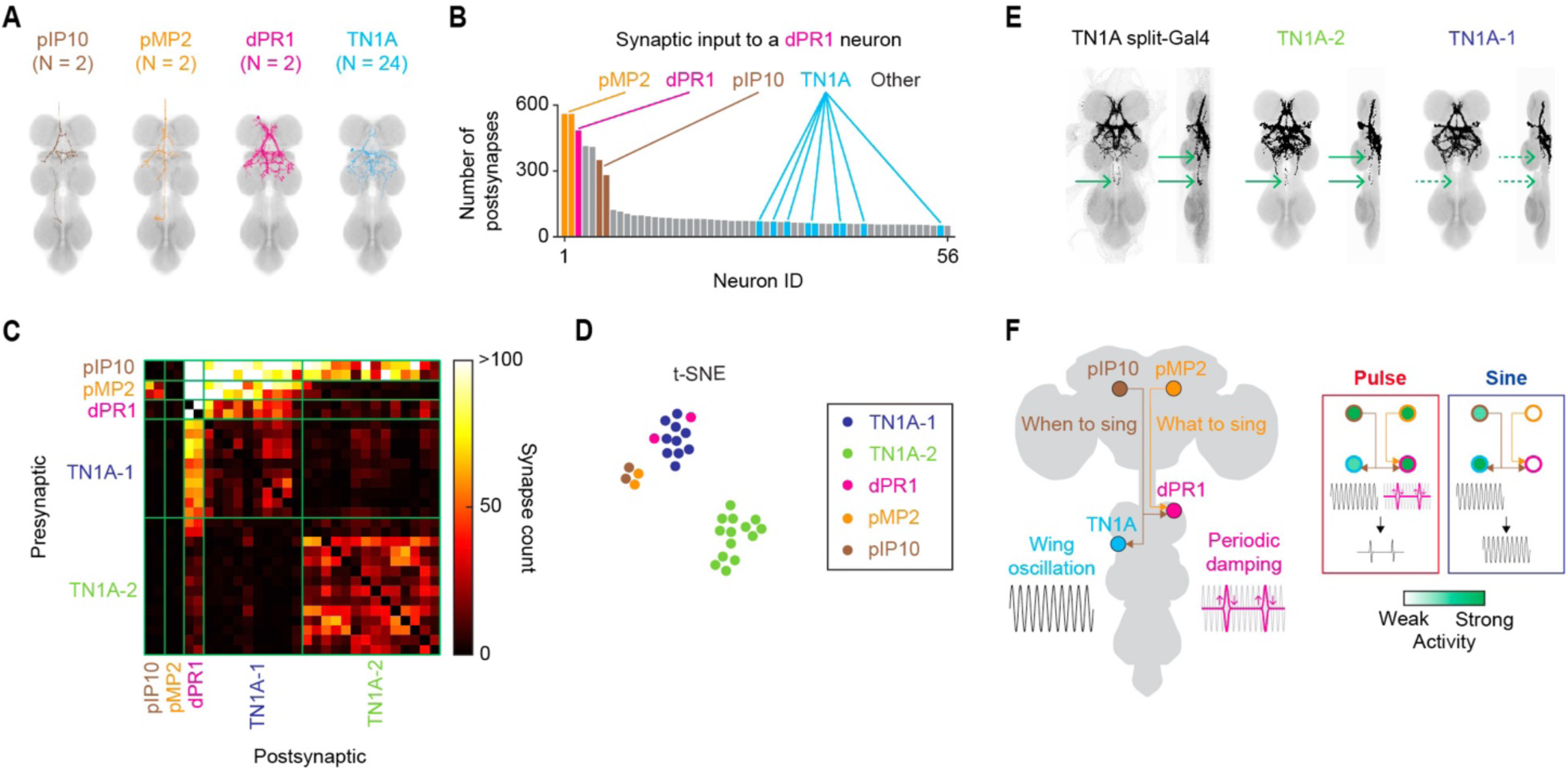
Neural circuit architecture for flexible song production. (A) Electron microscopy (EM) reconstruction images of single pIP10, pMP2, dPR1, and TN1A neurons. (B) Histogram of the number of synapses from upstream neurons onto a dPR1 neuron (body ID: 10300). Only neurons with 50 or more synapses onto dPR1 are shown. (C) Synaptic connectivity matrix among pIP10, pMP2, dPR1, and TN1A neurons. (D) A t-distributed stochastic neighbor embedding plot generated from the connectivity matrix in (C). (E) Expression pattern of the TN1A split-Gal4 (left) and the overlaid EM reconstruction images of TN1A-2 (middle) and TN1A-1 (right) neurons. Arrows indicate the neurites that were present in the TN1A split-Gal4 and TN1A-2 neuron images but not in TN1A-1 neuron images. (F) Model of flexible song production.

dPR1 and pMP2 showed similar activity patterns during song, raising the possibility that dPR1 receives input from pMP2. Indeed, pMP2 provides the largest numbers of input synapses to dPR1 (Figure 7B), suggesting that dPR1 derives its pulse-selectivity from pMP2. pIP10 also provides a substantial number of input synapses to dPR1 (Figure 7B), although pIP10 but not dPR1 was active during sine song (Figures 1 and 6). These observations suggest that dPR1R requires concurrent input from pIP10 and pMP2 for activation.

We next examined TN1A neurons and found that these neurons can be separated into two groups based on connectivity (Figures 7C and 7D). One group, which we term TN1A-1 (N = 10 neurons), receives a large number of synaptic inputs from pIP10, pMP2, and dPR1. On the other hand, the second group, TN1A-2 (N = 14 neurons), receives substantial input from pIP10, but not from pMP2 or dPR1. TN1A-1 neurons, but not TN1A-2 neurons, have a large number of synaptic connections to dPR1. Furthermore, connections within each group of TN1A neurons were stronger than those across the two TN1A groups. Overall, TN1A-1 and dPR1 had similar connectivity, whereas TN1A-2 neurons display connectivity distinct from other cell types (Figures 7C and 7D).

The TN1A split-Gal4 line we used for activity imaging labels approximately 8 neurons per fly, indicating that it includes a subset of TN1A neurons identified in the connectome dataset. The TN1A neurons labeled by the driver line showed activity patterns more similar to those of pIP10 than to dPR1 or pMP2, suggesting that the driver line targets primarily or entirely TN1A-2 neurons. Consistent with this idea, we found that the driver line labels neurites that were present only in the TN1A-2 population (Figure 7E). Together with the observations of neural activity, these results suggest that the combinatorial activity patterns produced in the VNC are inherited from descending pathways that encode information about when and what song to sing (Figure 7F).

## Discussion

Studies of motor circuits during naturalistic behavior, in both invertebrates and vertebrates, have often been hampered by the inaccessibility of the relevant circuits for observation in behaving animals. To overcome this limitation, we developed a novel two-photon calcium imaging assay for recording neural activity from the ventral nerve cord (VNC) of *Drosophila* males while they performed moment-to-moment switching between alternative courtship behaviors. We found that overlapping populations of VNC neurons are activated in a combinatorial manner when flies alternate between different types of courtship song. A population of song-related descending neurons also exhibit combinatorial activation patterns associated with song switching. Connectomic analysis revealed systematic synaptic connections between these descending and VNC neurons, consistent with a model in which parallel descending pathways drive overlapping yet distinct motor modules. These findings suggest that the *Drosophila* song system employs combinatorial coding to pattern action sequences during courtship.

Previous studies that involved manipulation of specific neurons in the VNC often observed changes in the amount of either pulse or sine song^21, 23^, leading to the proposal that different types of song are produced by independent neural populations^18, 21^. Contrary to this view, we found that although optogenetic stimulation of TN1A neurons induces sine song, these neurons are in fact active during both pulse and sine song. This observation is consistent with our recent finding that silencing the activity of TN1A neurons influences the amplitude of pulse song as well as the amount of sine song^23^. It was shown previously that a population of wing control muscles are active during both pulse and sine song, as we observe for TN1A neurons^10^, suggesting that TN1A neurons shape the pattern of activity in these muscles. These muscles have been proposed to control features of wing movement relevant for both pulse and sine song, such as the angle of attack^10, 11^. Indeed, silencing wing motor neurons for some of these muscles alters properties of both types of song^11^. Thus, pulse and sine song involve activation of a shared population of VNC neurons and wing muscles that may shape wing oscillations that are common to both song types.

In contrast to TN1A, dPR1 neurons are active during pulse but not sine song. Similarly, it was shown previously that a group of wing control muscles are active during pulse but not sine song^10^, suggesting that these muscles are controlled by dPR1 neurons. These muscles are thought to contribute to the damping of wing oscillations during inter-pulse intervals as well as the initiation of wing movement at the end of each inter-pulse interval^10, 11^. While some wing muscles are active during both song types and some are selective for pulse song, no wing muscle is specifically active during sine song^10, 11, 27^. This suggests that a group of muscles generate wing oscillations during both pulse and sine song production while additional muscles are recruited during pulse song to shape inter-pulse intervals. Our cell-type-specific and pan-neuronal recordings in the VNC show that this combinatorial recruitment pattern is present in the song motor system – some neurons are active during both types of song and most of the song-type selective neurons prefer pulse song. Connectome analysis suggests that members of the two neural populations, one not selective for song type and the other preferring pulse song, are reciprocally connected. These findings suggest that the song motor system contains two neural populations that independently operate as a functional module for a specific motor element, and the activation pattern of these modules determines which type of song is produced.

The neural population that prefers pulse song was mainly composed of neurons expressing *fru*. This result complements previous findings that *fru*-expressing neurons in the VNC are causally involved in pulse song production^19, 20, 23^. The VNC song circuit is capable of modulating specific pulse song parameters, such as the amplitude of song depending on the distance to females^15^, suggesting that individual *fru* neurons encode specific features of pulse song. Indeed, manipulating the activity of different *fru*-expressing neuron types affects different pulse song parameters^19, 23^. Similarly, silencing different wing motor neurons causes changes in different aspects of pulse song^11, 13^. Thus, the neural circuit for pulse song may be composed of subnetworks that control specific features of pulse song through activation of distinct muscles.

Courting males rapidly switch between pulse and sine song based on the behavioral context^7^. The activity of pulse-preferring neurons in the VNC, including dPR1, tracked ongoing song type, suggesting that these neurons play a key role in context-dependent song production. The pulse-specific activity of dPR1 is likely to be inherited from the descending neuron pMP2, which causally contributes to pulse song production^23^ and provides primary input to dPR1. We propose that pMP2 plays a role in controlling song type by regulating the activation states of the motor module that dPR1 belongs to. On the other hand, the descending neuron pIP10 was active during both pulse and sine song and provides input to both dPR1 and TN1A. Furthermore, pIP10 is causally involved in production of both pulse and sine song^16, 23^. These findings suggest that pIP10 determines the timing of singing without specifying song type. Thus, descending neurons innervating the song motor system may form parallel pathways that independently specify when to sing and what to sing.

Production of diverse movements by combinatorial activity patterns is not unique to the *Drosophila* song system. For example, in the mammalian respiratory system, a group of neurons are active during fictive normal breathing, whereas a subset of these active neurons become inactive during fictive gasping^42^. Notably, normal breathing and gasping are likely to involve activation of overlapping sets of muscles^3^, similar to the production of pulse and sine songs^10, 27^. In contrast, movements driven by independent sets of muscles, as in the case of alternating movements of left and right limbs during locomotion^2^, may be controlled through separate neural populations. Thus, the pattern of muscle recruitment required for distinct movements may bias evolution toward either independent or combinatorial neural control of muscles.

In summary, we uncovered the functional circuit architecture for flexible courtship song production in *D. melanogaster*. Our novel calcium imaging assay in courting flies complements existing strengths of this system such as cell type specific drivers for activity recording and manipulation^11, 16, 19, 21, 23^ and comprehensive circuit analysis with the connectome^41^. Together, the *Drosophila* song system will provide a unique opportunity to investigate the circuit mechanisms of neural computations underlying action selection and motor pattern generation.

## Acknowledgments

We thank A. Leonardo, V. Goncharov, and J. Arnold for help setting up the two-photon microscope, S. Sawtelle, B. Arthur, and E. Behrman for help with song recording, B. Arthur for help with SongExplorer, the Janelia Project Technical Resources team (C. Christoforou, K. Klose, Y. He, and J. Hausenfluck, led by G. Ihrke) and the FlyLight team for performing dissections, histological preparations, and confocal imaging in immunohistochemical experiments, Z. Dorman, G. W. Meissner, and G. Ihrke for help with multi-color flip out experiment designs, H. Otsuna for help with split-Gal4 line generation and analysis of the EM dataset, G. M. Rubin and V. Jayaraman for fly stocks, V. Ruta for comments on the manuscript, and members of the Stern lab for valuable discussions. This article is subject to HHMI’s Open Access to Publications policy. HHMI lab heads have previously granted a nonexclusive CC BY 4.0 license to the public and a sublicensable license to HHMI in their research articles. Pursuant to those licenses, the author-accepted manuscript of this article can be made freely available under a CC BY 4.0 license immediately upon publication.

## Author contributions

H.M.S. and D.L.S. conceived the study. H.M.S. performed all experiments and analyzed data. H.M.S., K.W., J.L.L., M.S., and B.J.D. generated genetic reagents. H.M.S. and D.L.S. wrote the manuscript with input from K.W., J.L.L., and B.J.D.

## Methods

### Fly husbandry

Flies (*Drosophila melanogaster*) were raised on cornmeal-molasses food at 23 °C under a 12-h/12-h light-dark cycle until eclosion. In all experiments, we used 5–7 days old adult male flies collected within 6 hours after eclosion and maintained in isolation in the dark. For the adult male flies used in experiments involving optogenetic stimulation, the food was supplemented with 0.4 mM all-trans-retinal (Toronto Research Chemicals). Virgin female flies used in behavioral experiments were housed in groups of approximately twenty.

### Fly stocks and genotypes

Fly stocks used in this study are as follows: w^1118^ (Stern Laboratory); UAS-CsChrimson-tdTomato(VK00005)^26^ (a gift from Gerald Rubin); VT037566-p65ADZp(attP40)^43, 44^ (Dickson Laboratory); VT041688-ZpGDBD(attP2)^44^ (Bloomington Drosophila Stock Center; BDSC #72408); VT042732-p65ADZp(attP40)^44^ (BDSC #71534); dsx-ZpGDBD^21^; UAS-CsChrimson-mVenus(attP18)^26^ (BDSC #55134); R22D03-LexA(attP2)^25^ (a gift from Gerald Rubin); LexAop2-CsChrimson-tdTomato(VK00005)^26^ (a gift from Vivek Jayaraman); UAS-IVS-jGCaMP7f(su(Hw)attP5)^33^ (BDSC #80906); UAS-IVS-myr::tdTomato(attP40)^25^ (BDSC #32222); w^1118^,hsFLPG5.PEST(attP3);;UAS-FRT.stop-myr::smGdP-HA(VK00005),UAS-FRT.stop-myr::smGdP-V5-THS-UAS-FRT.stop-myr::smGdP-FLAG(su(Hw)attP1)^45^ (BDSC #64085); VT040556-p65ADZp(attP40)^16, 44^ (BDSC #72060); VT040347-ZpGDBD(attP2)^16, 44^ (BDSC #75302); VT040347-ZpLexADBD(attP2)^44^ (Dickson Laboratory); UAS-IVS-jGCaMP7s(su(Hw)attP5)^33^ (BDSC #80905); dsx-Gal4^46^ (a gift from Bruce Baker); R57C10-LexA(attP40)^25^ (BDSC #52817); UAS-CsChrimson-tdTomato(su(Hw)attP5)^26^ (a gift from Vivek Jayaraman); LexAop2-IVS-jGCaMP7s(VK00005)^33^ (BDSC #80913); R57C10-Gal4(attP2)^47^ (BDSC #39171); fru-Gal4^39^ (BDSC #66696); fru-Flp^22^; tubP-FRT.stop-Gal80;MKRS/TM6B^48^ (BDSC #38878); R57C10-Gal4(attP40)^47^ (a gift from Gerald Rubin); VT026873-p65ADZp(attP40)^44^ (BDSC #86831);VT028160-ZpGdbd(attP2)^44^ (BDSC # 73842); 5xUAS-IVS-myr::smGFP-FLAG(VK00005)^49^ (a gift from Gerald Rubin); 3xUAS-Syt::smGFP-HA(su(Hw)attP1)^49^ (a gift from Gerald Rubin).

The dPR1, TN1A, and pMP2 split-Gal4 drivers were generated from the following split lines: dPR1 from SS65789 (VT037566-p65ADZp(attP40); VT041688-ZpGDBD(attP2)); TN1Afrom SS59832 (VT042732-p65ADZp(attP40);dsx-ZpGDBD/TM6b); pMP2 from SS43275 (VT026873-p65ADZp(attP40);VT028160-ZpGDBD(attP2)). These lines were identified by screening split lines identified by color depth MIP search^50^ of candidate *fru*-expressing neurons and neurons identified with trans-Tango^51^ applied to a pIP10 split-Gal4 line^16, 23^.

The genotypes of the flies used in each figure are as follows. Figure 1A, S1J, and S1N, w^1118^/Y;;UAS-CsChrimson-tdTomato(VK00005)/+; Figures 1B, 7E, S1A, S1C, S1D, S4G, and S4H, w^1118^,UAS-CsChrimson-mVenus(attP18)/Y;VT037566-p65ADZp(attP40)/+;VT041688-ZpGDBD(attP2)/+, w;VT042732-p65ADZp(attP40)/+;dsx-ZpGDBD/UAS-CsChrimson-tdTomato(VK00005); Figures 1D and 1E, S1G–J, VT037566-p65ADZp(attP40)/+;VT041688-ZpGDBD(attP2)/UAS-CsChrimson-tdTomato(VK00005), VT042732-p65ADZp(attP40)/+;dsx-ZpGDBD/UAS-CsChrimson-tdTomato(VK00005); Figures 1G and S1K, R22D03-LexA(attP2)/LexAop2-CsChrimson-tdTomato(VK00005); Figures 1H and 1L–O, UAS-IVS-jGCaMP7f(su(Hw)attP5),UAS-IVS-myr::tdTomato(attP40)/+;R22D03-LexA(attP2),LexAop2-CsChrimson-tdTomato(VK00005)/+; Figures 1I–M, S1R–X, S1FF, S3A, and S3B, UAS-IVS-jGCaMP7f(su(Hw)attP5),UAS-IVS-myr::tdTomato(attP40)/VT037566-p65ADZp(attP40);R22D03-LexA(attP2),LexAop2-CsChrimson-tdTomato(VK00005)/VT041688-ZpGDBD(attP2); Figures 1N–Q, S1Y–EE, S1GG, and S3B, UAS-IVS-jGCaMP7f(su(Hw)attP5),UAS-IVS-myr::tdTomato(attP40)/VT042732-p65ADZp(attP40);R22D03-LexA(attP2),LexAop2-CsChrimson-tdTomato(VK00005)/dsx-ZpGDBD; Figure S1B, w^1118^,hsFLPG5.PEST(attP3)/Y;VT037566-p65ADZp(attP40)/+;UAS-FRT.stop-myr::smGdP-HA(VK00005),UAS-FRT.stop-myr::smGdP-V5-THS-UAS-FRT.stop-myr::smGdP-FLAG(su(Hw)attP1)/VT041688-ZpGDBD(attP2); Figure S1E, w^1118^,hsFLPG5.PEST(attP3)/Y; VT042732-p65ADZp(attP40)/+;UAS-FRT.stop-myr::smGdP-HA(VK00005),UAS-FRT.stop-myr::smGdP-V5-THS-UAS-FRT.stop-myr::smGdP-FLAG(su(Hw)attP1)/dsx-ZpGDBD; Figures S1J, w^1118^/Y;VT037566-p65ADZp(attP40)/+;VT041688-ZpGDBD(attP2)/+, w^1118^/Y;VT042732-p65ADZp(attP40)/+;dsx-ZpGDBD/+; Figures S1P and S1Q, VT040556-p65ADZp(attP40)/+;VT040347-ZpGDBD(attP2)/UAS-CsChrimson-tdTomato(VK00005); Figures 2A and S2A, VT040556-p65ADZp(attP40),UAS-IVS-jGCaMP7s(su(Hw)attP5)/VT040556-p65ADZp(attP40);VT040347-ZpLexADBD(attP2),LexAop2-CsChrimson-tdTomato(VK00005)/VT040347-ZpGDBD(attP2); Figures 2B and S2B, VT040556-p65ADZp(attP40)/+;VT040347-ZpLexADBD(attP2)/LexAop2-CsChrimson-tdTomato(VK00005); Figures 2C–E, S2D, S2E, and S3E–J, VT040556-p65ADZp(attP40),UAS-IVS-jGCaMP7s(su(Hw)attP5)/+;VT040347-ZpLexADBD(attP2),LexAop2-CsChrimson-tdTomato(VK00005)/dsx-Gal4; Figures 2F–H, S2F, and S2G, VT040556-p65ADZp(attP40),UAS-IVS-jGCaMP7s(su(Hw)attP5)/VT042732-p65ADZp(attP40);VT040347-ZpLexADBD(attP2),LexAop2-CsChrimson-tdTomato(VK00005)/dsx-ZpGDBD; Figure S2C, w^1118^/Y;;LexAop2-CsChrimson-tdTomato(VK00005)/+, w^1118^/Y;VT040556-p65ADZp(attP40)/+;VT040347-ZpLexADBD(attP2)/+; Figure 3, S3C, and S3D, UAS-IVS-jGCaMP7f(su(Hw)attP5),UAS-IVS-myr::tdTomato(attP40)/+;R22D03-LexA(attP2),LexAop2-CsChrimson-tdTomato(VK00005)/dsx-Gal4; Figures 4C–H, 5J, 5L, and S4A–C, UAS-IVS-jGCaMP7f(su(Hw)attP5),UAS-IVS-myr::tdTomato(attP40)/+;R22D03-LexA(attP2),LexAop2-CsChrimson-tdTomato(VK00005)/R57C10-Gal4(attP2); Figures S4E and S4F, VT037566-p65ADZp(attP40),UAS-CsChrimson-tdTomato(su(Hw)attP5)/R57C10-LexA(attP40);VT041688-ZpGDBD(attP2),LexAop2-IVS-jGCaMP7s(VK00005)/+, VT042732-p65ADZp(attP40),UAS-CsChrimson-tdTomato(su(Hw)attP5)/R57C10-LexA(attP40);dsx-ZpGDBD/LexAop2-IVS-jGCaMP7s(VK00005); Figures 5B–F and S5, UAS-IVS-jGCaMP7f(su(Hw)attP5),UAS-IVS-myr::tdTomato(attP40)/+;R22D03-LexA(attP2),LexAop2-CsChrimson-tdTomato(VK00005)/fru-Gal4; Figures 5H–L, UAS-IVS-jGCaMP7f(su(Hw)attP5)/R57C10-Gal4(attP40),tubP-FRT.stop-Gal80;R22D03-LexA(attP2),LexAop2-CsChrimson-tdTomato(VK00005)/fru-Flp; Figures 6B–F, S6A, and S6B, UAS-IVS-jGCaMP7f(su(Hw)attP5),UAS-IVS-myr::tdTomato(attP40)/VT040556-p65ADZp(attP40);R22D03-LexA(attP2),LexAop2-CsChrimson-tdTomato(VK00005)/VT040347-ZpGDBD(attP2); Figures 6B, 6G–J, S6C, and S6D, UAS-IVS-jGCaMP7f(su(Hw)attP5),UAS-IVS-myr::tdTomato(attP40)/VT026873-p65ADZp(attP40);R22D03-LexA(attP2),LexAop2-CsChrimson-tdTomato(VK00005)/VT028160-ZpGDBD(attP2).

### Behavioral experiments

Song recordings in freely moving flies were conducted using the multi-channel song recording system described previously^52^. Briefly, the system is composed of 5-mm behavioral chambers, each equipped with a microphone (CMP5247TF-K, CUI Devices). The output of the microphones was recorded at 5 kHz. Temperature and relative humidity were maintained at 23°C and 50%, respectively. For optogenetic stimulation, an array of red LEDs (630-nm) (NFLS-R300X3-WHT-LC2, Super Bright LEDs) was placed approximately 15 mm from the tops of the chambers. During the recordings, the chambers were constantly illuminated with an ambient blue light (1.4 μW/mm^2^) (STN-BBLU-A3A-10B5M-12V, Super Bright LEDs) to enable flies to perform visually guided behaviors such as orientation toward females.

Flies were transferred from culture vials to the chambers with an aspirator without anesthesia. After one male fly was placed in each chamber, continuous light stimulation at a fixed power was applied for 10 s with an inter-trial interval of 20 s in each trial. Light power varied from trial-to-trial (6 levels between 4.0 and 39.7 μW/mm^2^) in a randomized order. Stimulation of each light power was repeated in 6 trials.

### Calcium imaging

Flies were prepared for calcium imaging in the VNC using the protocol described previously^53^ with modifications. Briefly, flies were placed on a Peltier plate that maintained temperature at 4 °C. The proboscis was fixed to the head capsule with UV glue, and the middle and hind legs were cut around the femur-trochanter junctions. The flies were then attached to a custom holding plate upside down (Figure 1F) by applying the glue to the coxa and trochanter of the middle legs and femur of the forelegs. The thorax was largely free from the glue so as not to disturb thorax oscillations and thus wing movement. The cuticles and apodemes covering the pro- and meso-thoracic ganglia were removed with fine forceps. The dissection was performed in saline (103 mM NaCl, 3 mM KCl, 5 mM N-tris(hydroxymethyl) methyl-2aminoethane-sulfonic acid, 8 mM trehalose, 10 mM glucose, 26 mM NaHCO_3_, 1 mM NaH_2_PO_4_, 1.5 mM CaCl_2_, and 4 mM MgCl_2_; pH, 7.1–7.3; osmolarity, 270-275 mOsm) bubbled with carbogen (95% O_2_ and 5% CO_2_). Calcium imaging in the brain was performed using the same protocol as described before^53^ except that legs were not removed.

Calcium imaging was conducted using a two-photon microscope (Bergamo II, Thorlabs) with a pulsed laser tuned to 940 nm (InSight X3, Spectra-Physics). The laser power measured under the objective lens was kept below 25 mW. A piezo actuator (PFM450E, Thorlabs) was used to move an objective lens (N16XLWD-PF, Nikon) for data acquisition from multiple z-planes. For experiments with optogenetic activation of pIP10, dPR1, or TN1A, two-dimensional images (512 × 512 pixels, pixel size 0.49 um) were taken from 30 z-planes (4 um step) at 1.4 volumes per s. For experiments with optogenetic activation of the song driver, images (256 × 256 pixels, pixel size 0.25–0.99 um) were taken from 10 z-planes (4 um step) at 7.1 volumes per s. In the recording from single dPR1 neurons, imaged volumes contained cell bodies of these neurons in both hemispheres in most of the experiments. In the recording from single TN1A and TN1 neurons, imaged volumes contained cell bodies of most of the target neurons in one hemisphere. In the recording from pIP10 and pMP2 neurons, imaged volumes contained the neurites of these neurons in a dorsal part of the brain. In a subset of recordings where neurons expressed tdTomato in addition to GCaMP, signals of both fluorescent proteins were recorded simultaneously. Carbogenated saline was perfused throughout the recording.

During calcium imaging in the VNC with song driver stimulation, optogenetic activation light (660-nm) (S1FC660, Thorlabs) passed through a long-pass filter (FEL0650, Thorlabs) and was delivered with a patch cable (M125L01, Thorlabs) placed at 6 mm away from the fly. The long-pass filter was not used in other calcium imaging experiments. For each fly, a continuous light stimulation at a fixed power was applied for 10 s with an inter-trial interval of 20 s. The light power varied from trial-to-trial (6 levels, 4.9–156.2 μW/mm^2^ for pIP10, dPR1, and TN1A stimulations and recording in the brain, 4.9–17.1 μW/mm^2^ for song driver stimulation during recording in the VNC) in a randomized order. Stimulation of the same power was repeated in 6 trials for song driver stimulation and 3 trials for pIP10 stimulation. For the pIP10 and song driver simulations in tethered flies without calcium imaging, the light power ranged from 2.4 to 14.6 μW/mm^2^. The same stimulus was presented in 6 trials.

The sound of vibrating wings was recorded with a pair of microphones (NR-23158, Knowles) placed near the left and right wings, respectively. Microphone signals were amplified with a custom amplifier^52^ and recorded at 10 kHz. Wing movement was monitored with a camera (BFS-U3-04S2M-CS, Teledyne FLIR) equipped with a periscopic lens (InfiniStix 90°, working distance 94 mm, magnification 1.0×, Infinity). Flies were illuminated by infrared light (850-nm) (M850F2, Thorlabs) with a patch cable (M125L01, Thorlabs). A long-pass (FEL0800, Thorlabs) and a short-pass (FES0850, Thorlabs) filter were placed on the lens to remove the optogenetic and two-photon activation light, respectively. Video was recorded at 100 frames per s.

### Immunohistochemistry

The dissections, immunohistochemistry, and imaging of fly central nervous systems were performed using protocols described previously^45, 49^ (https://www.janelia.org/project-team/flylight/protocols) with modifications. Briefly, brains and VNCs were dissected in Schneider’s insect medium and fixed in 2% paraformaldehyde at room temperature for 55 min. Tissues were washed in PBT (0.5% Triton X-100 in phosphate buffered saline) and blocked using 5% normal goat serum before incubation with antibodies.

To visualize the expression pattern of GFP or GCaMP, tissues were stained using rabbit anti-GFP (1:1000, A-11122, Thermo Fisher Scientific) and mouse nc82 (1:30, Developmental Studies Hybridoma Bank) as primary antibodies and Alexa Fluor 488-conjugated goat anti-rabbit (A-11034, Thermo Fisher Scientific) and Alexa Fluor 568-conjugated goat anti-mouse (A-11031, Thermo Fisher Scientific) as secondary antibodies. To visualize the expression pattern of tdTomato, tissues were stained using rabbit anti-dsRed (1:1000, #632496, Clontech) and mouse nc82 (1:30) as primary antibodies and Cy3-conjugated goat anti-rabbit (#111-165-144, Jackson ImmunoResearch) and Cy2-conjugated goat anti-mouse (#115-225-166, Jackson ImmunoResearch) as secondary antibodies. To visualize the expression pattern of GCaMP and tdTomato in the same samples, tissues were stained using chicken anti-GFP (1:1000, A-10262, Thermo Fisher Scientific), rabbit anti-dsRed (1:1000), and mouse nc82 (1:30) as primary antibodies and Alexa Fluor 488-conjugated goat anti-chicken (A32931, Thermo Fisher Scientific), Cy3-conjugated goat anti-rabbit, and Cy5-conjugated goat anti-mouse (#115-175-166, Jackson ImmunoResearch) as secondary antibodies. To visualize the expression pattern of *dsx*, tissues were stained using mouse anti-DsxDBD (1:2, Developmental Studies Hybridoma Bank), rabbit anti-GFP (1:1000), and rat anti-DN-Cadherin (1:100, DN-Ex #8, Developmental Studies Hybridoma Bank) as primary antibodies and Cy3-conjugated goat anti-mouse (#115-165-166, Jackson ImmunoResearch), Alexa Fluor 488-conjugated goat anti-rabbit, and Cy5-conjugated goat anti-rat (#112-175-167, Jackson ImmunoResearch) as secondary antibodies. Flies for multi-color flip out^45^ underwent a 40-min heat shock during 0–1 day post eclosion and dissected at 5–14 days old. These tissues were stained with the antibodies described previously^45^. The antibodies used for visualizing the expression pattern of pMP2 split-Gal4 line were described previously^49^.

Image stacks were collected using a confocal microscope (LSM710, Zeiss, Germany) with an objective lens (Plan-Apochromat 20x/0.8 M27 or Plan-Apochromat 40X/1.3 M27, Zeiss, Germany).

### Song type classification

Pulse and sine songs were detected using SongExplorer^54^, a deep-learning based algorithm for segmenting acoustic communication signals. We first removed low-frequency components of the microphone signals (<100 Hz), which were mostly irrelevant for song, using continuous wavelet transformation. We then performed manual annotation for a subset of the audio data. For the data obtained from freely moving flies, we used the labels “pulse”, “sine”, “inter-pulse intervals”, “others” for non-song sounds (e.g., noise during grooming), and “ambient.” Another label “flight” was added for the data from tethered flies because these flies occasionally showed flight-like wing movement. We then trained a classifier separately for freely moving, tethered flies during the recording from the VNC, and tethered flies during the recording from the brain using the annotated data. These classifiers were applied to assign a label at every 1.6 ms of the data from all the recordings. The classification accuracies, measured with cross-validation, for the data from freely moving and tethered flies were 91.6% and 88.1%, respectively. Visual inspection of the classifier predictions revealed that misclassifications often occurred at a smaller number of consecutive time bins compared with correct predictions. To correct for these misclassifications, we smoothed the time series of classifier predictions with median filters (the window size for detecting “pulse”, 17.6 ms; “sine”, 25.6 ms; “flight”, 80 ms).

To compare the amounts of pulse and sine song, we calculated the pulse train based on the classifier predictions. Pulse song is composed of discrete pulse events separated by inter-pulse intervals, whereas sine song is continuous. We defined the pulse train by merging pulse events with inter-pulse intervals of 50 ms or shorter. This allows comparisons of the amounts of pulse and sine song on the same scale. Events designated as “pulse” in the figures represent the pulse train.

### Data analysis for behavioral experiments

For each recording, we calculated the time series of the proportions of pulse and sine song by combining data across repeated presentations of the same optogenetic stimulus. These time series were averaged across flies to obtain the time course of song proportions, or across time to calculate the song probabilities for each fly. To quantify how much optogenetic stimulation changed the amounts of pulse and sine song, we subtracted the proportions of pulse and sine song during optogenetic stimulation from those during 5-s pre-stimulation periods.

### Data analysis for calcium imaging experiments

Time series of z-stack images underwent rigid motion correction with the NoRMCorre algorithm^55^. To analyze calcium signals of dPR1 cell bodies, we first set a volume of interest (VOI) separately for the dPR1 neuron in each hemisphere. We then calculated an average intensity z-projection of the VOI and performed another round of rigid motion correction. The same procedure was carried out for analyzing TN1A cell bodies, dPR1 neurite, TN1A neurite, pIP10 neurite, and pMP2 neurite, except that a single volume of interest was used to calculate an average intensity z-projection. For each recording, regions of interest (ROIs), each corresponding to a cell body or target neurites, were manually defined using an average image of the motion corrected z-projections. Each ROI for pIP10 neurite contained signals from the pIP10 neurons in both hemispheres. Each ROI for pMP2 neurite reflected signals from one pMP2 neuron. Signals in each ROI were averaged for further analysis. Motion corrections were performed using either GCaMP or tdTomato signals. For analyzing data from pan-neuronal and *fru*-expressing neuron imaging, the motion corrected z-stack images went through 2 by 2 voxel binning in the xy-planes to increase the signal-to-noise ratio. ΔF/F was calculated for each ROI or voxel by defining the baseline as the mean fluorescent signal during 10-s periods preceding optogenetic stimulation.

To compare spatial activity patterns across flies, inter-fly image registration was performed for the data from pan-neuronal and *fru*-expressing neuron imaging. First, the motion corrected z-stack images were averaged across time for each recording. Second, the averaged z-stack images went through 2 by 2 voxel binning in the xy-planes. Third, the binned images from one recording were selected as a reference for each target volume (e.g., a dorsal volume) and each driver line (e.g., the pan-neuronal line). Finally, the time-averaged z-stack images of each recording were aligned to the reference with either the NoRMCorre rigid registration or the Computational Morphometry Toolkit^56^ run with a Fiji plugin (https://github.com/jefferis/fiji-cmtk-gui). These registration parameters were used to calculate population averaged calcium response patterns.

To analyze the relationship between calcium signals and song, we converted the sampling rate of song classifier predictions (625 Hz) to that of calcium imaging (7.1 Hz; sampling at every 141 ms) as follows. Time series of the predictions were separated into 141-ms bins, and the mean probabilities of each event (“pulse”, “sine”, “ambient”, and “flight) were calculated for each bin. We considered that an event occurred in a bin if the probability exceeded a threshold (0.1 for “pulse”, 0.6 for “sine, 0.9 for “ambient”, 0.01 for “flight”). Defining event occurrence with these thresholds rarely gave false positives when assessed with visual inspection. Flight-like behavior induced changes in calcium signals in a large fraction of the imaged volume (data not shown), which can obscure song-related calcium signals. We therefore excluded from further analysis the data during “flight” and from -1 to 1 s after “flight”.

To characterize the changes in calcium signals during song-type transitions, we first calculated the average ΔF/F in the two bins over which song type was changed (from 141 ms before the transition to 141 ms after the transition). We then subtracted this value from the average ΔF/F in the third and fourth bins after the change in song type (282–564 ms after the transition) for the analysis of cell body signals. To take into account faster GCaMP kinetics at neurite than cell bodies^34^, the average ΔF/F in the second and third bins after the change in song type (141–423 ms after the transition) was used for the subtraction in the analysis of neurite signals. Considering the kinetics of jGCaMP7f^33^, this quantity likely reflects changes in neural activity during song type transitions. We averaged this quantity across transition events and divided it by the average ΔF/F in the two bins over which song type was changed, obtaining the normalized mean change in ΔF/F during song type transitions for each ROI. To characterize the changes in calcium signals during quiet to pulse transitions, we detected “pulse” bins that followed at least 2-s continuous “ambient” periods. The changes in ΔF/F during the transitions were defined as the subtraction of the average ΔF/F during the 2-s period before the transition from that after the transition. We analyzed the data from the recordings where the transitions occurred at least five times for the analysis of song type transitions and two times for the analysis of quiet to pulse transitions. We focused on song type transitions rather than calcium signals during each song type because pulse and sine song tended to occur in different timings in 10-s stimulation periods, and thus factors irrelevant for song (e.g., time-varying input from song irrelevant CsChrimson expressing neurons) can contribute to the differential calcium signals during pulse and sine song.

To assess whether individual TN1 neurons responded to optogenetic stimulation, we calculated the mean ΔF/F during the stimulation period subtracted from that of the 10-s pre-stimulation period for each trial for a range of stimulation levels (9.8–17.1 μW/mm^2^ for song driver stimulation; 19.5–156.2 μW/mm^2^ for pIP10 stimulation). We then aggregated the responses across stimulation levels and performed one-sample t-test. We performed the same test was performed for dPR1, and TN1A neurons and found that all these neurons displayed significant responses (p < 0.05). We also tested pIP10 and pMP2 with a range of stimulation levels (230.2–575.6 μW/mm^2^) and found that all but two pMP2 neurons showed significant responses (p < 0.05). These two pMP2 neurons were excluded from further analysis.

For single dPR1, TN1A, and TN1 neurons, the preference for song type was quantified using the receiver operating characteristics (ROC) analysis (Figures S3A). We first built the distribution of the changes in ΔF/F separately for pulse-to-sine and sine-to-pulse transitions. We then used the two distributions to calculate the area under the ROC curve. An index termed the song-type preference was defined as the AUC scaled from −1 to 1. This index quantifies how much the type of transitions (pulse-to-sine vs. sine-to-pulse) can be predicted based on the changes in calcium signals. A song-type preference of 1 and -1 represents perfect predictions with higher calcium signals for pulse and sine song, respectively. A song-type preference of 0 indicates that the changes in calcium signals were indistinguishable between the two types of transitions. For single TN1A neurons, we assessed whether the song-type preference was different from 0 using a permutation test. We shuffled the labels (i.e., pulse-to-sine and sine-to-pulse transitions) of transition responses and calculated song-type preference. This was repeated 10,000 times to obtain a null distribution of song-type preference. The same permutation test was conducted for each voxel for the data obtained in pan-neuronal and *fru*-expressing neuron imaging (Figures 4 and 5). For these data, we also calculated a null distribution of transition responses for each voxel using permutation (Figures 4 and 5). We first randomized the relationship between song type transitions and calcium signals by shuffling the trials for calcium signal data. We then calculated the distribution of transition responses as in the original data.

Images involved in calcium signals were spatially smoothed with a 2-D Gaussian filter for presentation in figures.

### Connectomic and morphological analysis

All connectomic and neuron skeleton data were obtained from the male adult nerve cord dataset^41^ using neuPrint^57^. Only the neurons that have been traced and possess at least 100 presynapses of postsynapses were analyzed. EM-reconstructed images of single neurons were created by registering neuron skeletons to a VNC template made from light microcopy images^58^. To create the t-distributed stochastic neighbor embedding plot from the connectivity matrix among pIP10, pMP2, dPR1, and TN1A neurons (Figure 7D), we took each neuron and constructed a vector that represents the number of input synapses to this neuron from each neuron and the number of output synapses from this neuron to each of the other neurons. We then performed t-SNE using the cosine distance metric with the perplexity of 10. A list of body IDs for the neurons analyzed was provided in Table S1.

## Statistics

All statistical tests were two-tailed and performed using MATLAB 2022b.

## Data availability

Data are available at FigShare.

## Code availability

Code is available at FigShare.

## Supplementary information

**Video S1**

An example trial of calcium imaging in dPR1 neurons. Top left: a bottom view of the fly. The left side of the image corresponds to the right side of the fly. Bottom left: the microphone signal for the right wing. Top right: raw fluorescent signals (F) averaged across z-planes. The left side of the image corresponds to the left side of the VNC. Top left: schematic of the imaged volume. Red boxes represent the timing of optogenetic stimulation.

**Video S2**

An example trial of calcium imaging in TN1A neurons. Same format as in Video S1.

**Video S3**

An example trial of pan-neuronal calcium imaging in dPR1 neurons. Top left: the microphone signal for the right wing. The red box represents the timing of optogenetic stimulation. Bottom left: schematic of the imaged volume. Right: raw fluorescent signals (F) for each z-plane.

**Video S4**

Same as in Video S3 but for calcium signals averaged across z-planes.

---

**Figure S1.**
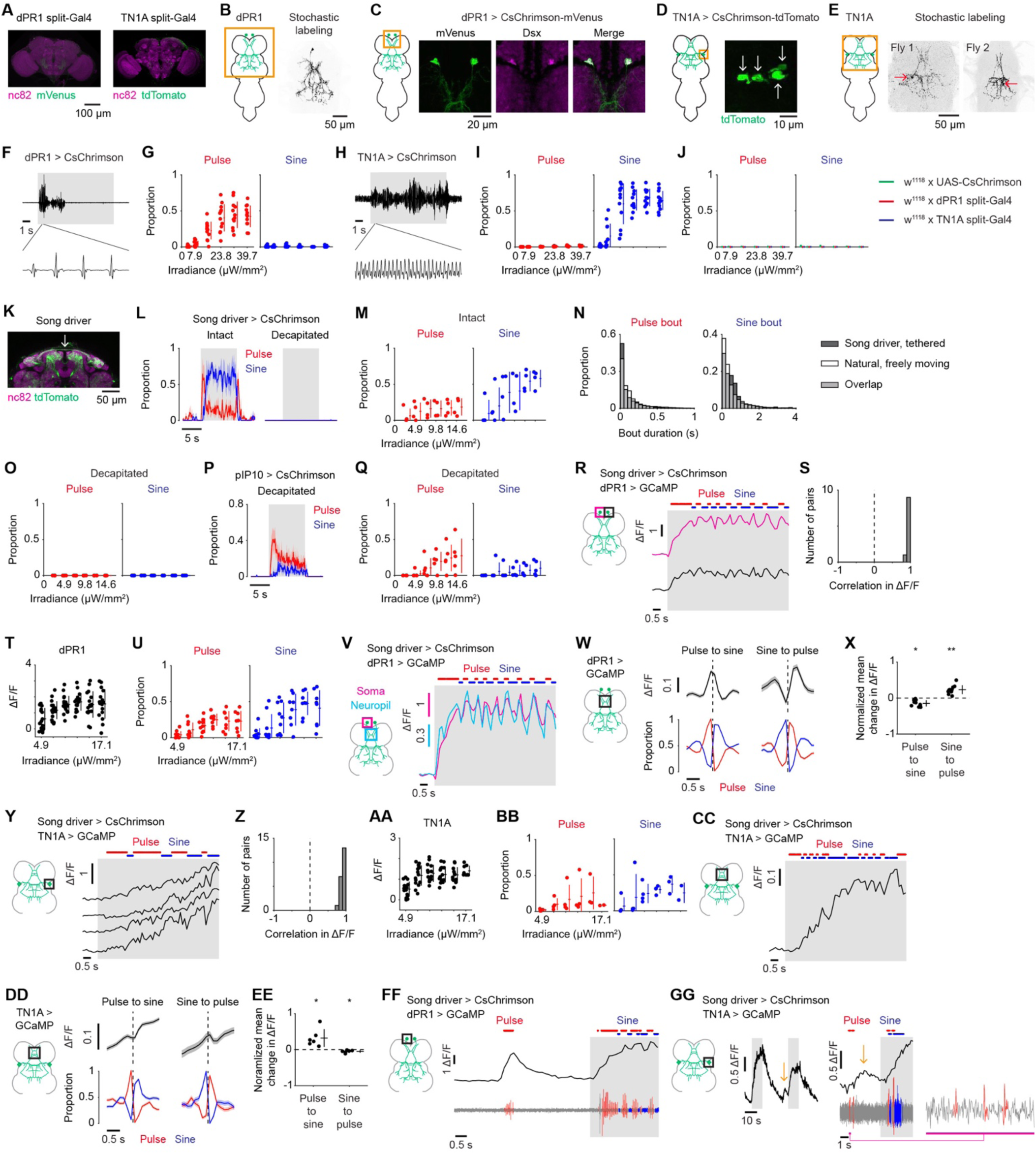
Flexible song production involves combinatorial activation of dPR1 and TN1A neurons. (A) Expression patterns of the dPR1 split-Gal4 (left) and the TN1A split-Gal4 (right) in the brain. (B) A single neuron visualized in the VNC with stochastic labeling using the dPR1 split-Gal4 line. (C) *Dsx* is expressed in the neurons labeled with the dPR1 split-Gal4 line. (D) Cell body expression pattern of CsChrimson in the TN1A split-Gal4 line. Arrows indicate the location of cell bodies. (E) Examples of single TN1A neurons visualized with stochastic labeling using the TN1A split-Gal4. Arrows indicate the cell bodies. (F) Example of the sound induced by optogenetic activation of dPR1. The shaded area represents the period where LED stimulation was applied. (G) The mean proportions of pulse (left) and sine (right) song during the stimulation period. Each dot represents a fly. Lines represent mean ± SD across flies (N = 12 flies). (H and I) Same as (F) and (G) but for TN1A (N = 12 flies). (J) Results for control flies in the optogenetic activation experiment. The mean proportions of pulse and sine song during the stimulation period. Lines represent mean ± SD across flies (N = 11 flies for UAS-CsChrimson; N = 12 flies for dPR1 split-Gal4; N = 12 flies for TN1A split-Gal4). (K) Expression pattern of the song driver at an anterior-dorsal part of the central brain of a male fly. The arrow indicates the commissural fibers of P1 neurons. (L) Flies expressing CsChrimson under the control of the song driver were attached to the recording plate and optogenetically stimulated with a red laser. For the decapitated group, the head was removed after attachment to the recording plate. No thoracic dissection was made on these flies. The graphs show the time course of the proportions of pulse and sine song during laser stimulation at an irradiance of 12.2 μW/mm^2^. Data are represented as mean ± SEM across flies (N = 4 flies for the intact group; N = 6 flies for the decapitated group). (M) The mean proportions of pulse (left) and sine (right) song during the stimulation period for the intact group. Each dot represents a fly. Lines represent mean ± SD across flies (N = 4 flies). (N) The distributions of the durations of pulse and sine song bouts. Dark grey bars, the songs induced by optogenetic activation with the song driver (N = 4 intact flies). White bars, the songs in a group of control flies in the optogenetic activation experiment shown in Figure S1J (*w*^1118^ crossed to UAS-CsChrimson; N = 11 flies). (O) Same as (M) but for the decapitated group (N = 6 flies). (P) Flies expressing CsChrimson under the control of the split-Gal4 line for pIP10 were attached to the recording plate, decapitated and provided with optogenetic stimulation. No dissection was made on the thorax. Graphs show time courses of the proportions of pulse and sine song during laser stimulation at an irradiance of 12.2 μW/mm^2^. Data are represented as mean ± SEM across flies (N = 5 flies). (Q) Same as (O) but for pIP10 (N = 5 flies). (R) Example ΔF/F traces of two dPR1 neurons recorded simultaneously. The top trace is the same as the one shown in Figure 1J. The song driver was used to express CsChrimson. The shaded area represents the period where laser stimulation was applied. (S) Histogram of the Pearson’s correlation coefficients between pairs of dPR1 neurons recorded simultaneously (N = 10 pairs in 10 flies). (T) The mean ΔF/F during the stimulation period for dPR1. Each dot represents a neuron. Lines represent mean ± SD across neurons (N = 16‒20 neurons in 8–10 flies for each irradiance). (U) The mean proportions of pulse and sine song during the stimulation period in the recordings from dPR1. Each dot represents a fly. Lines represent mean ± SD across flies (N = 8–10 flies for each irradiance). (V) Example ΔF/F trace of a part of the dPR1 neurite (cyan) along with that of a cell body recorded simultaneously (magenta). The trace for the cell body is the same as the one shown in Fig. 1J. (W) Time course of dPR1 neurite ΔF/F and song around song-type transitions. Dashed vertical lines represent the transition times. Data are from 10 flies and represented as mean ± SEM across transitions for both ΔF/F and song (N = 1092 and 429 events for pulse-to-sine and sine-to-pulse transitions, respectively). (X) The mean change in ΔF/F after song-type transitions relative to ΔF/F before the transitions for the neurite of dPR1. Each dot represents a fly. Lines represent mean ± SD across flies (N = 10 flies). *P < 0.05/2, **P < 0.01/2, one-sample t test with Bonferroni correction, the denominator represents the number of comparisons. (Y) Example ΔF/F traces of four TN1A neurons recorded simultaneously. The top trace is the same as the one shown in Figure 1N. (Z) Histogram of the Pearson’s correlation coefficients between pairs of TN1A neurons recorded simultaneously (N = 21 pairs in 4 flies). (AA) Same as (T) but for TN1A (N = 8–15 neurons in 2–4 flies). (BB) Same as (U) but for the recording from TN1A (N = 2–4 flies for each irradiance). (CC) Example ΔF/F trace of a part of the TN1A neurite. (DD and EE) Same as (W) and (X) but for TN1A. (DD) N = 318 pulse-to-sine and 122 sine-to-pulse transitions in 6 flies. (EE) N = 6 flies. (FF) Example ΔF/F trace of a single dPR1 neuron (top) together with the simultaneously recorded sound (bottom). The fly produced a train of pulse song outside the stimulation period, which was accompanied by an increase in calcium signals. (GG) Example ΔF/F trace of a single TN1A neuron. The fly produced pulse song outside the stimulation period, which was followed by an increase in calcium signals (arrows).

**Figure S2.**
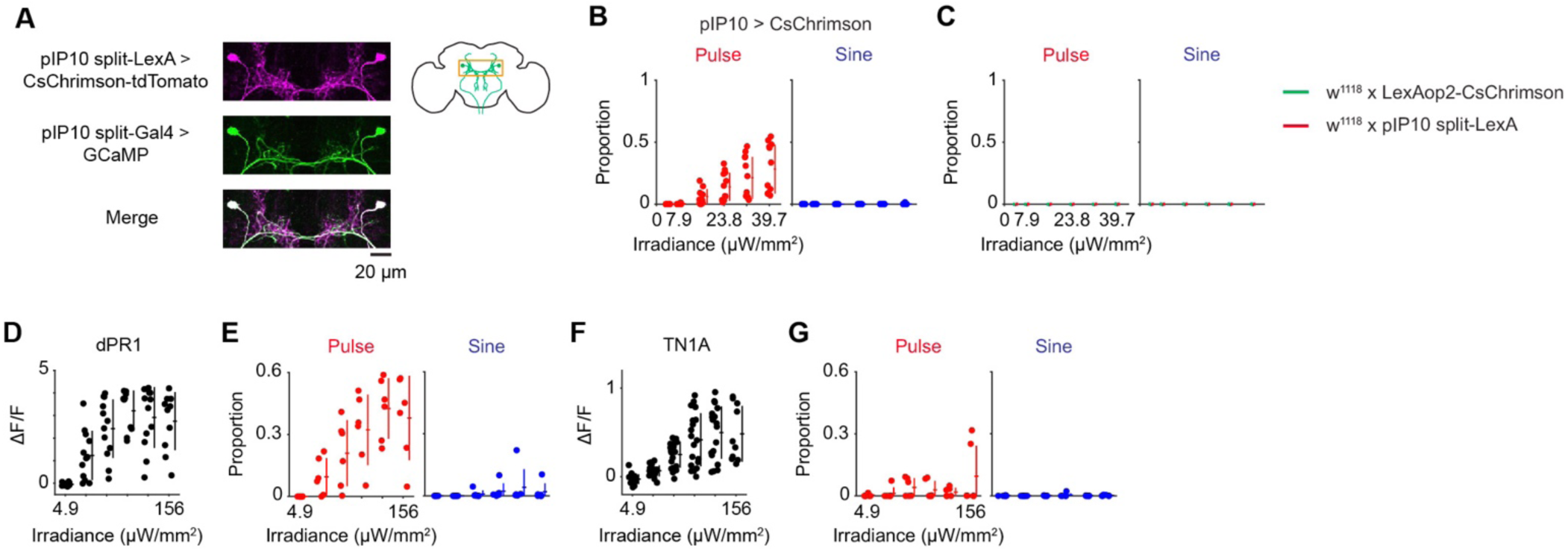
dPR1 and TN1A are activated during pulse song induced by stimulation of the descending neurons pIP10. (A) Expression patterns of the pIP10 split-LexA and split-Gal4 drivers around the cell bodies of pIP10. (B) The mean proportions of pulse and sine song during the stimulation period in the optogenetic activation experiment. Each dot represents a fly. Lines represent mean ± SD across flies (N = 12 flies). (C) Results for control flies in the optogenetic activation experiment. The mean proportions of pulse and sine song during the stimulation period.. Lines represent mean ± SD across flies (N = 12 flies for each genotype). (D) The mean ΔF/F of dPR1 during the stimulation of pIP10. jGCaMP7s was expressed with *dsx*-Gal4 and calcium signals were recorded from dPR1 cell bodies. Each dot represents a neuron. Lines represent mean ± SD across neurons (N = 12 neurons in 6 flies). The X-axis is in a logarithmic scale. (E) The mean proportions of pulse and sine song during the stimulation period in the recordings from dPR1. Each dot represents a fly. Lines represent mean ± SD across flies (N = 6 flies). (F and G) Same as (D) and (E) but for TN1A. (F) N = 19 neurons in 6 flies. (G) N = 6 flies.

**Figure S3.**
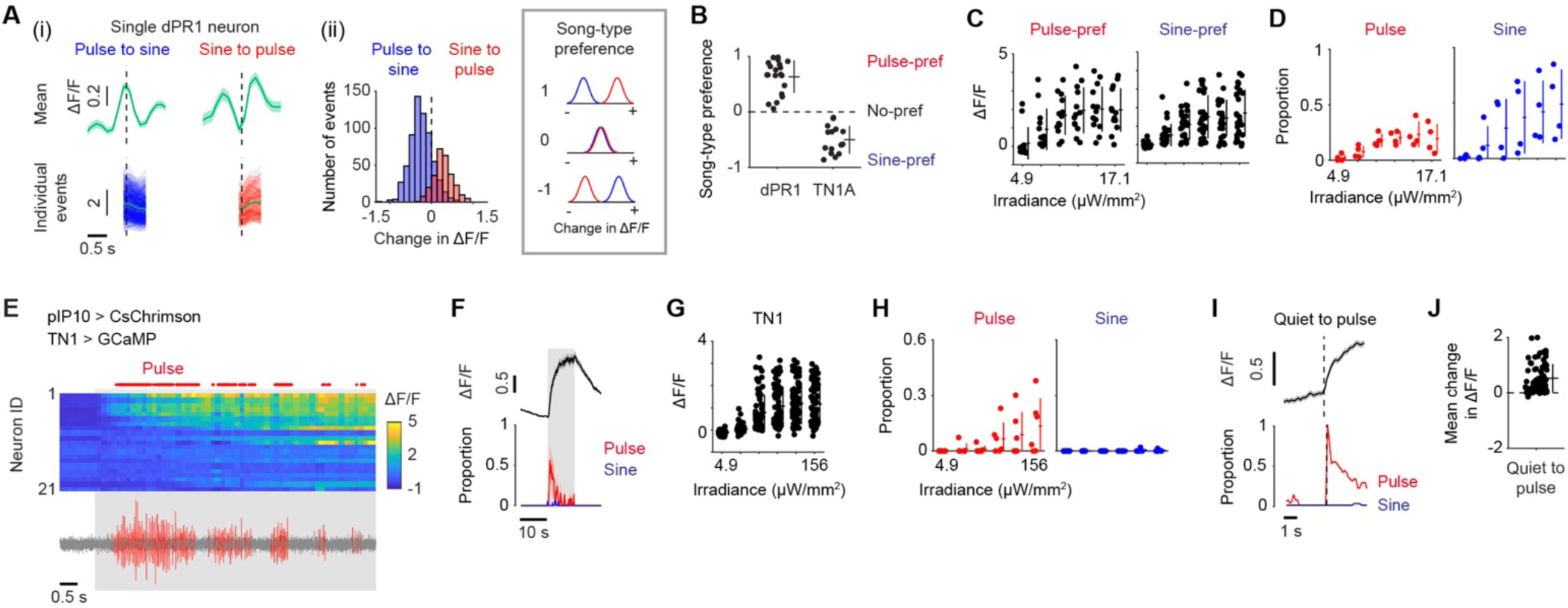
TN1 neurons exhibit combinatorial activation during flexible song production. (A) Calculation of the song-type preference for an example dPR1 neuron. This index quantifies the selectivity for pulse and sine song during song-type transitions for individual neurons and is calculated in two steps (see STAR Methods for detail). (i) First, the time course of ΔF/F around the transition is calculated for pulse-to-sine and sine-to-pulse transitions. As an example, the mean ΔF/F across events (top) and individual ΔF/F traces for each event (bottom) around song-type transitions are shown for one dPR1 neuron. Dashed vertical lines represent the transition time. For individual traces, only the part of the data used in the calculation of the index is shown. (ii) Second, histograms are calculated for the change in ΔF/F around the song-type transitions for pulse-to-sine (blue) and sine-to-pulse (red) transitions and the song-type preference is calculated using the receiver operating characteristics analysis. The song-type preference for this example dPR1 neuron was 0.87. (B) Histograms for the song-type preference for dPR1 (N = 20 neurons in 10 flies) and TN1A (N = 15 neurons in 4 flies) calculated using the data shown in Figure 1. (C) The mean ΔF/F for pulse-preferring and sine-preferring TN1 neurons during the stimulation of the neurons labeled by the song driver. Each dot represents a neuron. Lines represent mean ± SD across neurons (N = 13 pulse-preferring and 25 sine-preferring neurons in 4 flies for each irradiance). (D) The mean proportions of pulse and sine song during the stimulation period for the data shown in (C) (N = 2–4 flies for each irradiance). Each dot represents a fly. (E–J) Calcium imaging from TN1 cell bodies during simulation of pIP10. CsChrimson was expressed using the pIP10 split-LexA driver. (E) Example ΔF/F of a population of TN1 neurons recorded simultaneously (top) together with the sound recording (bottom). The shaded area represents the period where laser stimulation was applied. (F) Top, Time course of ΔF/F for the TN1 neurons that showed significant responses to optogenetic stimulation (p < 0.05, t-test; N = 77 neurons in 4 flies) for the laser stimulation at the irradiance of 78.1 μW/mm^2^. Bottom, The proportions of song during the recording from these neurons. Data are represented as mean ± SEM across neurons (top) and flies (bottom). (G) The mean ΔF/F for pulse-preferring and sine-preferring TN1 neurons during stimulation of pIP10. Each dot represents a neuron. Lines represent mean ± SD across neurons (N = 77 neurons in 4 flies). The X-axis is in a logarithmic scale. (H) The mean proportions of pulse and sine song during the stimulation period for the data shown in (G) (N = 4 flies). Each dot represents a fly. (I) Time course of TN1 ΔF/F and song around transitions from quiet to pulse song production. Dashed vertical lines represent the transition times. Data are from 77 neurons in 4 flies and represented as mean ± SEM across transitions for both ΔF/F and song (N = 869 transitions). (J) The mean change in ΔF/F after quiet-to-pulse transitions (see STAR Methods for detail) for TN1. Each dot represents a neuron. Lines represent mean ± SD across neurons (N = 77 neurons in 4 flies).

**Figure S4.**
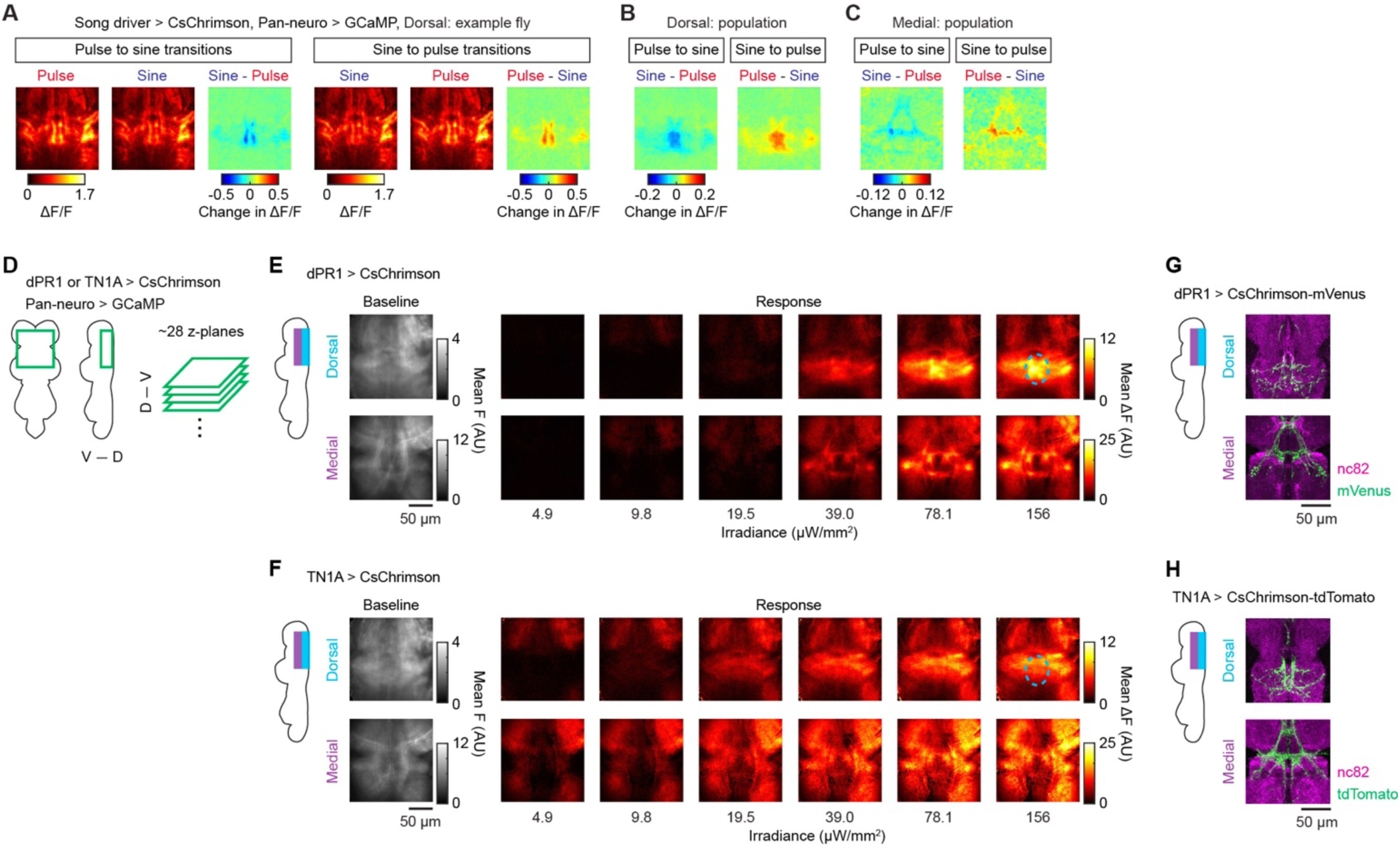
Combinatorial activation is a general feature in the song motor circuit. (A) Changes in calcium signals around song-type transitions for the example fly shown in Figure 5D. Left, average frames during pulse to sine transitions (N = 71 transitions). Images were averaged across z-planes and time. Right, average frames during sine to pulse transitions (N = 29 transitions). (B) Population averaged changes in ΔF/F during song-type transitions for the recordings from the dorsal part of the VNC (N = 9 flies). (C) Same as (B) but for the recordings from the medial part (N = 3 flies). (D) Schematic of the VNC region imaged. GCaMP was expressed using a pan-neuronal driver. CsChrimson was expressed with either the dPR1 or TN1A driver. (E) Pan-neuronal calcium imaging of the wing-related neuropils while activating dPR1. Frames were averaged across trials and flies (N = 6 trials for each irradiance; N = 7 flies), followed by further averaging across z-planes separately for dorsal and medial volumes of the VNC. The dashed circle in the “Dorsal” image for the irradiance of 156 μW/mm^2^ highlights a region that showed strong activation during dPR1 but not TN1A stimulation. (F) Same as (E) but for TN1A (N = 6 trials for each irradiance; N = 6 flies). (G) dPR1 neurons in the dorsal and medial volumes visualized with the dPR1 split-Gal4 line. (H) Same as (G) but for TN1A neurons visualized with the TN1A split-Gal4 line.

**Figure S5.**
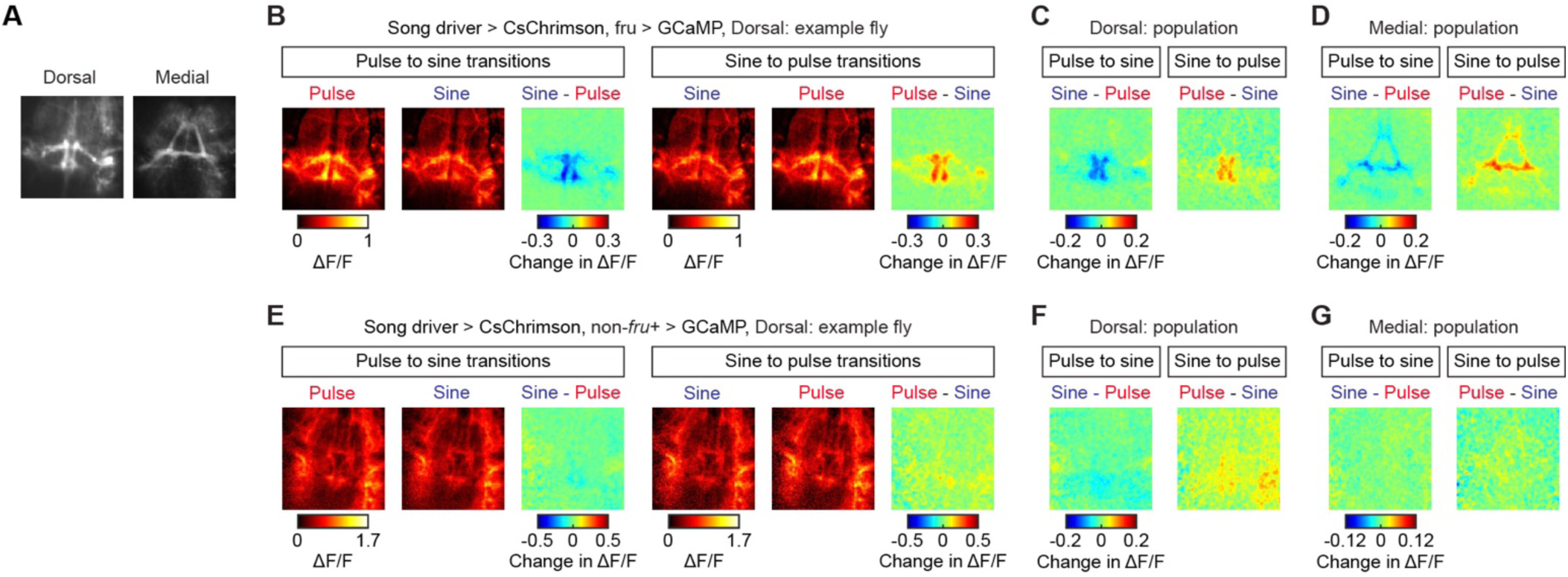
Song type selective signals in the circuit are carried by a genetically defined population of neurons.. (A) Example images of the calcium signals recoded from the dorsal (left) and medial (right) parts of the VNC. Images represent averages of frames across z-planes and time. (B) Changes in calcium signals across song-type transitions for the example fly shown in Figure 5B. Images were averaged across z-planes and time. Left, average frames during pulse to sine transitions (N = 137 transitions). Right, average frames during sine to pulse transitions (N = 65 transitions). (C) Population averaged changes in ΔF/F during song-type transitions for the recordings from the dorsal part of the VNC for *fru*-expressing neurons (N = 6 flies). (D) Same as (C) but for the recordings from the medial part (N = 5 flies). (E) Same as (B) but for non-*fru*-expressing neurons (N = 34 pulse to sine transitions, N = 13 sine to pulse transitions). (F and G) Same as (C) and (D) but for non-*fru*-expressing neurons. (F) N = 5 flies. (G) N = 3 flies.

**Figure S6.**
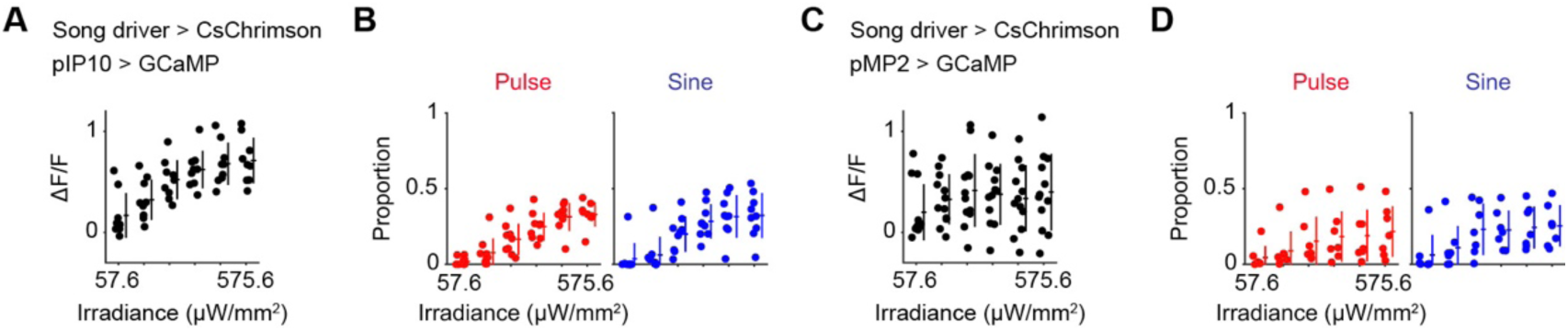
Flexible song production involves combinatorial activation of parallel descending pathways. (A) The mean ΔF/F of pIP10 during the stimulation of the song driver. Each dot represents a fly from which the combined calcium signals of the left and right pIP10 neurons were recorded. Lines represent mean ± SD across flies (N = 9 flies). The X-axis is in a logarithmic scale. (B) The mean proportions of pulse and sine song during the stimulation period in the recordings from pIP10. Each dot represents a fly. Lines represent mean ± SD across flies (N = 9 flies). (C and D) Same as (A) and (B) but for pMP2 except that the activity of individual neurons was analyzed. (C) N = 12 neurons in 7 flies. (D) N = 7 flies.

